# RadiSeq: a single- and bulk-cell whole-genome DNA sequencing simulator for radiation-damaged cell models

**DOI:** 10.1101/2025.02.08.637266

**Authors:** Felix Mathew, Luc Galarneau, John Kildea

**Affiliations:** McGill University

## Abstract

**Objective:** To build and validate a simulation framework to perform single-cell and bulk-cell whole genome sequencing simulation of radiation-exposed Monte Carlo cell models to assist radiation genomics studies.

**Approach:** Sequencing the genomes of radiation-damaged cells can provide useful insight into radiation action for radiobiology research. However, carrying out post-irradiation sequencing experiments can often be challenging, expensive, and time-consuming. Although computational simulations have the potential to provide solutions to these experimental challenges, and aid in designing optimal experiments, the absence of tools currently limits such application. Monte Carlo toolkits exist to simulate radiation exposures of cell models but there are no tools to simulate single- and bulk-cell sequencing of cell models containing radiation-damaged DNA. Therefore, we aimed to develop a Monte Carlo simulation framework to address this gap by designing a tool capable of simulating sequencing processes for radiation-damaged cells.

**Main Results:** We developed RadiSeq – a multi-threaded whole-genome DNA sequencing simulator written in C++. RadiSeq can be used to simulate Illumina sequencing of radiation-damaged cell models produced by Monte Carlo simulations. RadiSeq has been validated through comparative analysis, where simulated data were matched against experimentally obtained data, demonstrating reasonable agreement between the two. Additionally, it comes with numerous features designed to closely resemble actual whole-genome sequencing. RadiSeq is also highly customizable with a single input parameter file.

**Significance:** RadiSeq enables the research community to perform complex simulations of radiation-exposed DNA sequencing, supporting the optimization, planning, and validation of costly and time-intensive radiation biology experiments. This framework provides a powerful tool for advancing radiation genomics research.

## Introduction

Genome sequencing studies have provided valuable insights into the biophysical mechanisms of ionizing radiation (IR) mutagenesis (Youk et al. 2024; Adewoye et al. 2015). However, the high cost and technical challenges associated with these types of experiments, particularly single-cell whole-genome sequencing (ScWGS), often limit their feasibility (Gawad, Koh, and Quake 2016). To minimize the cost and uncertainty associated with post-irradiation genome sequencing studies, careful experimental design is crucial. To this end, we have developed a Monte Carlo (MC) simulation framework designed to simulate DNA sequencing of IR-damaged cells. This framework can be used to optimize experimental parameters and predict experimental outcomes, thereby reducing the risk of wasted resources and ensuring more efficient and informative experiments.

MC simulations are extensively used to model IR interactions probabilistically. Geant4 (Agostinelli et al. 2003) and its user-friendly wrapper TOPAS (Faddegon et al. 2020), PHITS (Niita et al. 2006), FLUKA (Ferrari et al. 2005), and EGSnrc (National Research Council of Canada. Metrology Research Centre. Ionizing Radiation Standards 2021) are some examples among the numerous MC simulation toolkits used in radiotherapy-related research. With the goal of advancing the science and understanding of radiobiological effects at the cellular and subcellular levels, some of these toolkits have been extended with track structure modelling to simulate IR interactions at the DNA scale; Geant4-DNA (Incerti et al. 2010), TOPAS-nBio (Schuemann, McNamara, Ramos-Méndez, et al. 2019), and PARTRAC (Friedland et al. 2011) are some examples. Various geometric nuclear DNA models have also been developed and are used in MC-based radiobiology investigations (McNamara et al. 2018; Sakata et al. 2020; Bernal et al. 2013; Bernal and Liendo 2009; Bertolet et al. 2022; Montgomery et al. 2021). With the abundance of MC toolkits and DNA models now available, it is possible to simulate in detail the interaction of IR in cells to obtain a complete description of the DNA damages introduced. A recently-published data standard, called the Standard DNA Damage (SDD) format, has also been developed to report the DNA damage information from such simulations (Schuemann, McNamara, Warmenhoven, et al. 2019).

Genome sequencing has matured into a valuable biological technique with numerous applications, including in the field of IR research. Various sequencing approaches and bioinformatic techniques have been employed to investigate IR-induced genomic effects including mutation signatures (Behjati et al. 2016; Kageyama et al. 2021), DNA damage (Murray, Hardie, and Gautam 2019), and genomic alterations (Nguyen et al. 2016; Youk et al. 2024). While these approaches have typically focused on using tumors or clonal populations of cells in which all cells harbor the same mutations, we hypothesized that ScWGS of heterogeneous IR-damaged cell samples could provide additional insights into IR-induced genomic alterations. Our novel experiments yielded promising results (Mathew et al. 2023). However, we found that conducting novel post-irradiation ScWGS experiments is complex, expensive, and time-consuming. In this context, we postulated that the ability to simulate ScWGS and other sequencing approaches in silico can help with experimental design and can serve as a valuable complement to the experiments themselves. However, despite our ability to model IR-induced DNA damage, we were faced with a lack of tools to simulate the sequencing of damaged DNA.

A plethora of tools have emerged for simulating next-generation sequencing (NGS) techniques, offering various models for different sequencing technologies and protocols. Notable recent reviews by Alosaimi et al. (2020), Escalona et al. (2016), and Zhao et al. (2017) have assessed many simulators, evaluating their strengths and weaknesses. While most simulators can reasonably imitate real-world NGS procedures, there is no single-best solution suitable for all experimental scenarios (Milhaven and Pfeifer 2023). Notably, none of the existing tools can simulate the sequencing of diploid cells with strand breaks, which is a requisite for modelling the sequencing of IR-damaged cells since IR will most likely create somatic heterozygous mutations. Thus, the development of a new sequencing simulation framework, specifically to integrate with MC-generated IR-damaged cell models, was a prerequisite for us to use sequencing simulations to guide our irradiation and ScWGS experiments.

Here, we present RadiSeq, an open-source simulation framework (Mathew and Kildea 2024) that we developed for Illumina (Illumina Inc. San Diego, California, USA) ScWGS and bulk-cell whole-genome sequencing (BcWGS) of IR-damaged MC-generated cell models that describe their DNA damage in the SDD format.

## Materials and Methods

RadiSeq was developed in C++ with a high degree of end-user customizability in mind. Figure 1 presents a schematic of RadiSeq’s processing pipeline. The main goal of developing RadiSeq was to emulate real experiments and predict sequencing outcomes. This stands in contrast to the purpose of most existing sequencing simulation tools that aim to produce “gold standard” sequencing data for the purposes of validating and assessing bioinformatics analysis pipelines. Consequently, RadiSeq was designed to allow users to customize the simulation by using an input parameter file that contains all of the adjustable parameters needed by the models provided in the simulation framework.

**Figure 1:**
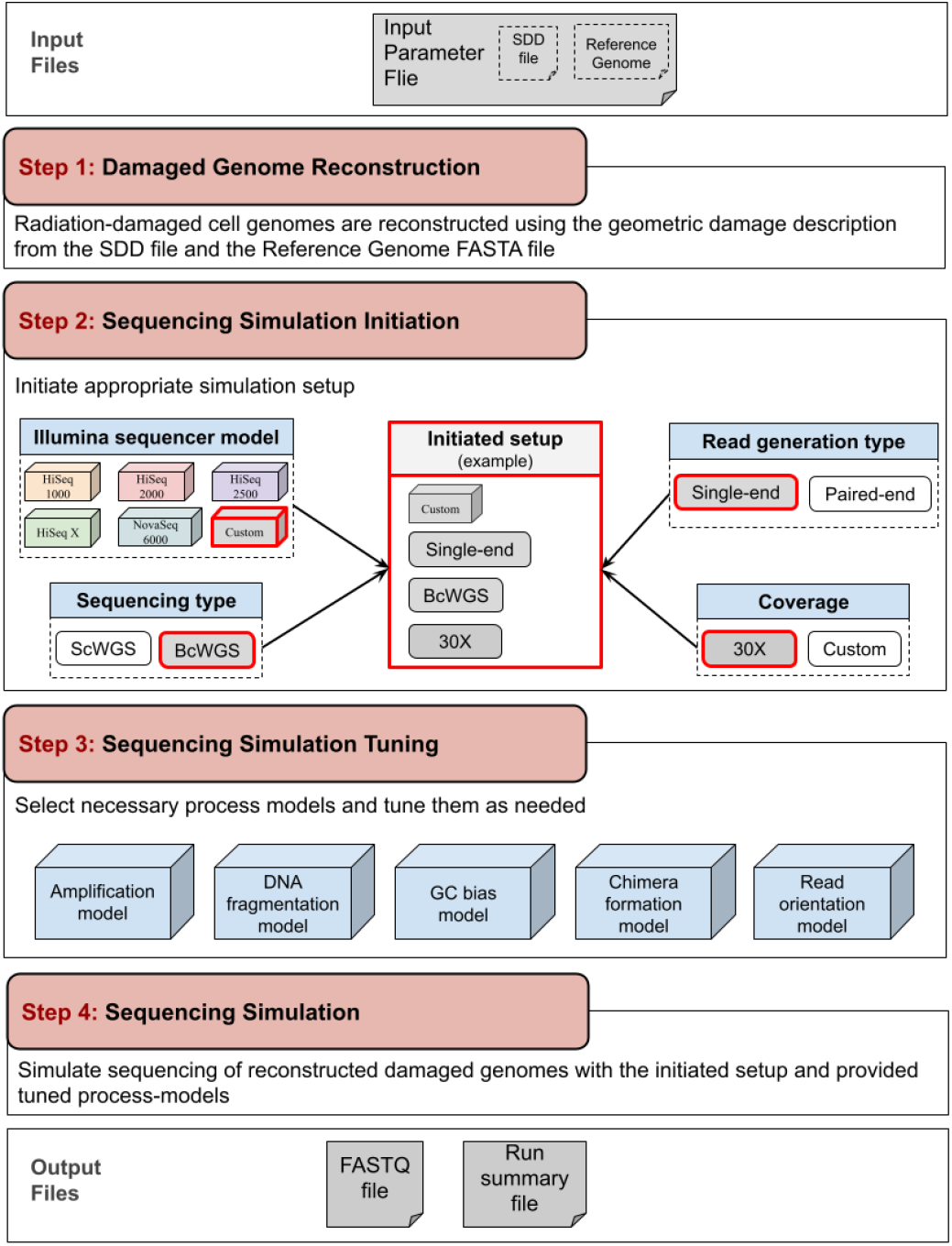
A schematic representation of RadiSeq’s sequencing simulation pipeline showing the four steps in a simulation. The red outline is used to delineate an example use case in Step 2.

### Step 1: Damaged Genome Reconstruction

RadiSeq’s ability to simulate the sequencing of IR-damaged DNA is what sets it apart from other sequencing simulators. DNA models used in MC irradiation simulations are geometric only–they record the location of DNA damage but contain no genomic sequence information. Therefore, RadiSeq’s first task is to create a sequenceable damaged genome. To do so, it takes two files as input, one containing the damage-annotated geometric genome of one or more cells in SDD format, as outputted by a MC pipeline, and the other containing a complete reference genome in the FASTA format (Lipman and Pearson 1985). As illustrated in Figure 2, a simple mapping algorithm then folds the reference genome onto the geometric genome, while maintaining the damage location annotations (base damages and strand breaks); the reference genome of any diploid organism, not necessarily human, can be used for this purpose.

**Figure 2:**
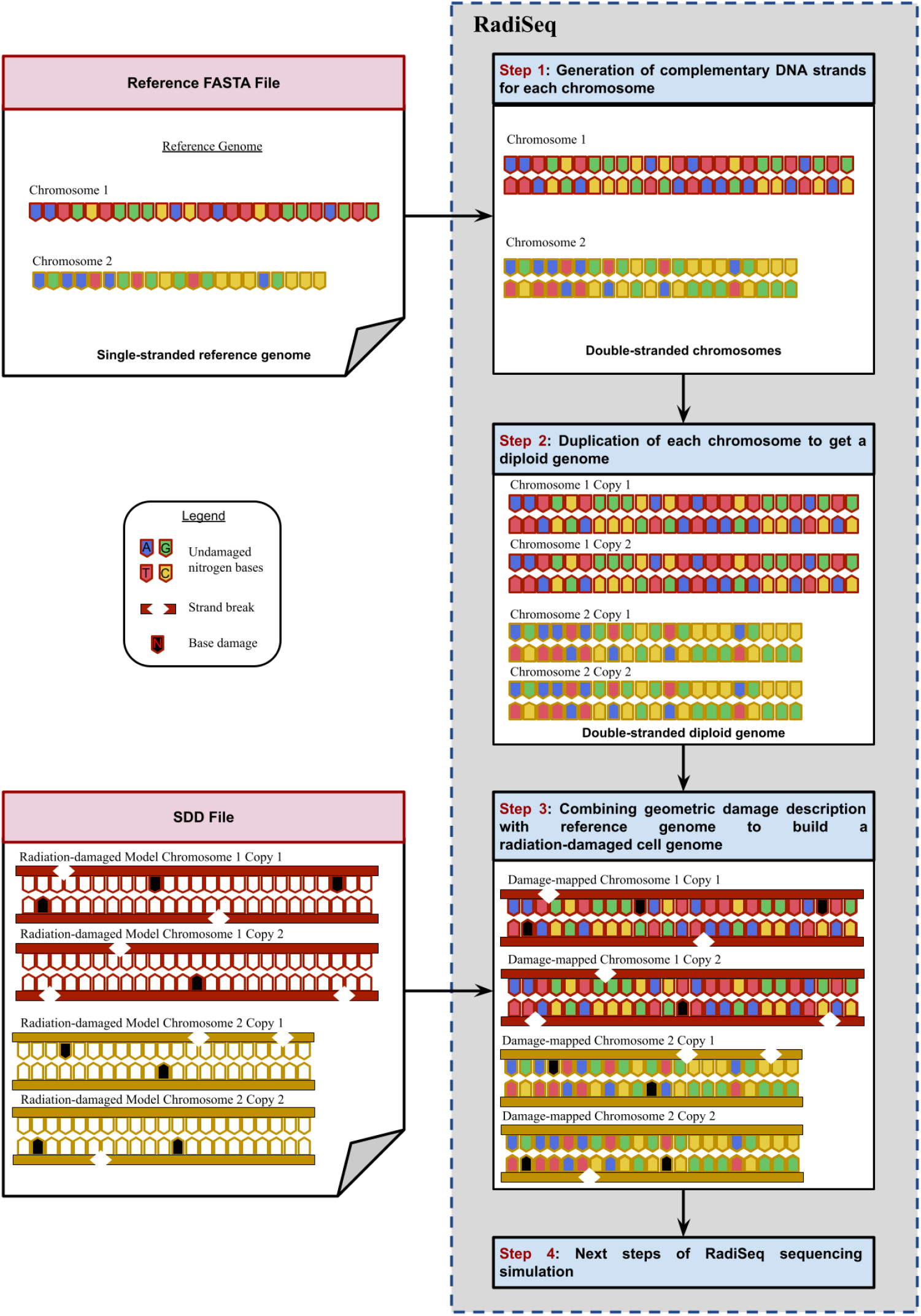
A schematic showing the steps of damaged-genome reconstruction in RadiSeq.

### Step 2: Sequencing Simulation Initiation

Given the variability in error-rate profiles and overall read quality across sequencing platforms, RadiSeq includes five pre-configured Illumina sequencer models and offers an option to specify a custom sequencer to include additional Illumina sequencers using user-provided error profiles. A sequencer profiling tool is bundled with RadiSeq to help users generate error profiles (see Supplementary Materials section 3.3). Users can choose between ScWGS and BcWGS to be performed on the reconstructed damaged genome. RadiSeq initiates the handling of reconstructed damaged genomes differently depending on this choice. Both single-end (SE) and paired-end (PE) read generation are supported.

### Step 3: Sequencing Simulation Tuning

To account for additional variability in PE read generation, RadiSeq includes options for different read-pair orientations such as forward-forward, reverse-forward, and others (see Supplementary Material). RadiSeq includes two DNA amplification models: (i) a uniform amplification model and (ii) a non-uniform multiple-displacement amplification (MDA) (Lasken 2009) model. RadiSeq’s modular design allows for easy expansion with additional models in the future. GC bias (Chen et al. 2013) is modeled using a unimodal triangular function. Additionally, chimeric read artifact formation (Lu et al. 2023) can be included in the simulation. All these pre-configured process models are easily customized and can be tuned to meet specific user needs.

### Step 4: Read Generation

The read-generation process differs between the ScWGS and BcWGS arms of RadiSeq. In BcWGS, each read or read pair is generated from a randomly-selected reconstructed cell within the sample, while in ScWGS, cells are processed sequentially. The desired coverage determines the total number of reads or the number of reads required per cell for BcWGS and ScWGS respectively. To construct a read (the first read in a pair for PE sequencing), RadiSeq evaluates the likelihood of a read originating from various genomic locations, considering factors such as GC bias and genome amplification. For PE reads, the fragmentation model is also employed to identify the probabilistic location of the second read in the pair. Read sequences, referred to as template sequences in RadiSeq, are then generated according to the specified read length and orientation. Finally, the generated template sequences are probabilistically modified by introducing chimeric sequences, insertions, deletions, and substitution errors, and their quality is adjusted accordingly to obtain the final reads. A detailed description of read generation is included in the supplementary materials (section: RadiSeq Simulation Procedure).

### Implementation and Performance

The generated read data are output as raw text files that follow the compressed FASTQ format specification (.fastq.gz) along with a RadiSeq run summary text file. A detailed schematic of the RadiSeq processing pipeline and additional details on each of the built-in models are included in the Supplementary Material.

Multithreading is enabled in RadiSeq and users can set the number of CPU threads for faster processing using the OpenMP parallel programming model (Chandra 2001), and operations are further optimized using memory mapping techniques. The current release of RadiSeq (RadiSeq_v2.0) requires C++ 17 or above, 64-bit (x64) CPU architecture, and is currently only compatible with Unix-based systems.

### Performance evaluation of RadiSeq

We have been using a human geometric nuclear DNA model that was built in-house and published open source as the ‘NICE model’ (Montgomery and Manalad 2022) for radiation exposure simulations in TOPAS-nBIO. To evaluate the performance of RadiSeq, the NICE model, which represents a human lymphocyte cell in the G0/1 phase, was sham-irradiated (i.e. no actual irradiation) to generate an undamaged SDD file. This file was used to independently simulate both ScWGS and BcWGS in RadiSeq because the simulation pipeline differs for each. During these sequencing simulations, CPU run time was measured to assess performance. Illumina PE reads of length 151 bp were successfully generated for varying genomic coverages using the NovaSeq 6000 sequencer (Illumina Inc.)–one of the models included in RadiSeq. The DNA fragment size distribution was specified using a beta function with a beta value of 10, a mode value of 155 bp, and a maximum of 200 bp. These settings were designed to closely match the conditions of the sequencing experiments previously performed in our lab. Other important parameters used in the simulation that may influence runtime included zero degree of GC bias, a uniform amplification model, and a read artifacts rate of 0.3. Simulations were performed with 40 CPU threads on an Intel Xeon Gold 6148 @ 2.40GHz processor of the Digital Research Alliance of Canada clusters.

### Verification and Validation of RadiSeq

Unit testing and integration testing were performed to ensure that the different components and models of RadiSeq functioned as expected. However, for the acceptance of RadiSeq, it was imperative to validate that the simulated data would accurately mimic the experimental data for both the BcWGS and ScWGS pipelines separately. For this purpose, we used aligned and analyzed experimentally obtained read data from sham-irradiated human B-lymphoblastoid cells in our lab. These cells were subjected to BcWGS and ScWGS experiments, and the corresponding experimental conditions were used in RadiSeq to generate simulated data for comparison. For ScWGS, one cell was randomly chosen from the experimental dataset to be used for this comparison.

The accuracy of RadiSeq was examined using the sham-irradiated NICE cell model. For the ScWGS, the MDA model was used to amplify single-cell genomes with a DNA fragment size distribution comparable to our ScWGS experiments (see Supplementary Methods). For the BcWGS simulation pipeline, the uniform amplification model and a DNA fragment size distribution comparable to our BcWGS experiments were used in the simulation. For both the ScWGS and BcWGS pipelines, RadiSeq-generated reads were analyzed using the FastQC tool (http://www.bioinformatics.babraham.ac.uk/projects/fastqc/) and then aligned to the reference human genome using the Bowtie 2 (Langmead and Salzberg 2012) aligner. Alignment statistics were obtained using Samtools (Li et al. 2009) and compared against the experimentally obtained data.

## Results

Figure 3 shows the RadiSeq computational performance in terms of CPU run time for both BcWGS and ScWGS simulations plotted on a semi-log graph. Linear fits to the data are also included for reference.

**Figure 3:**
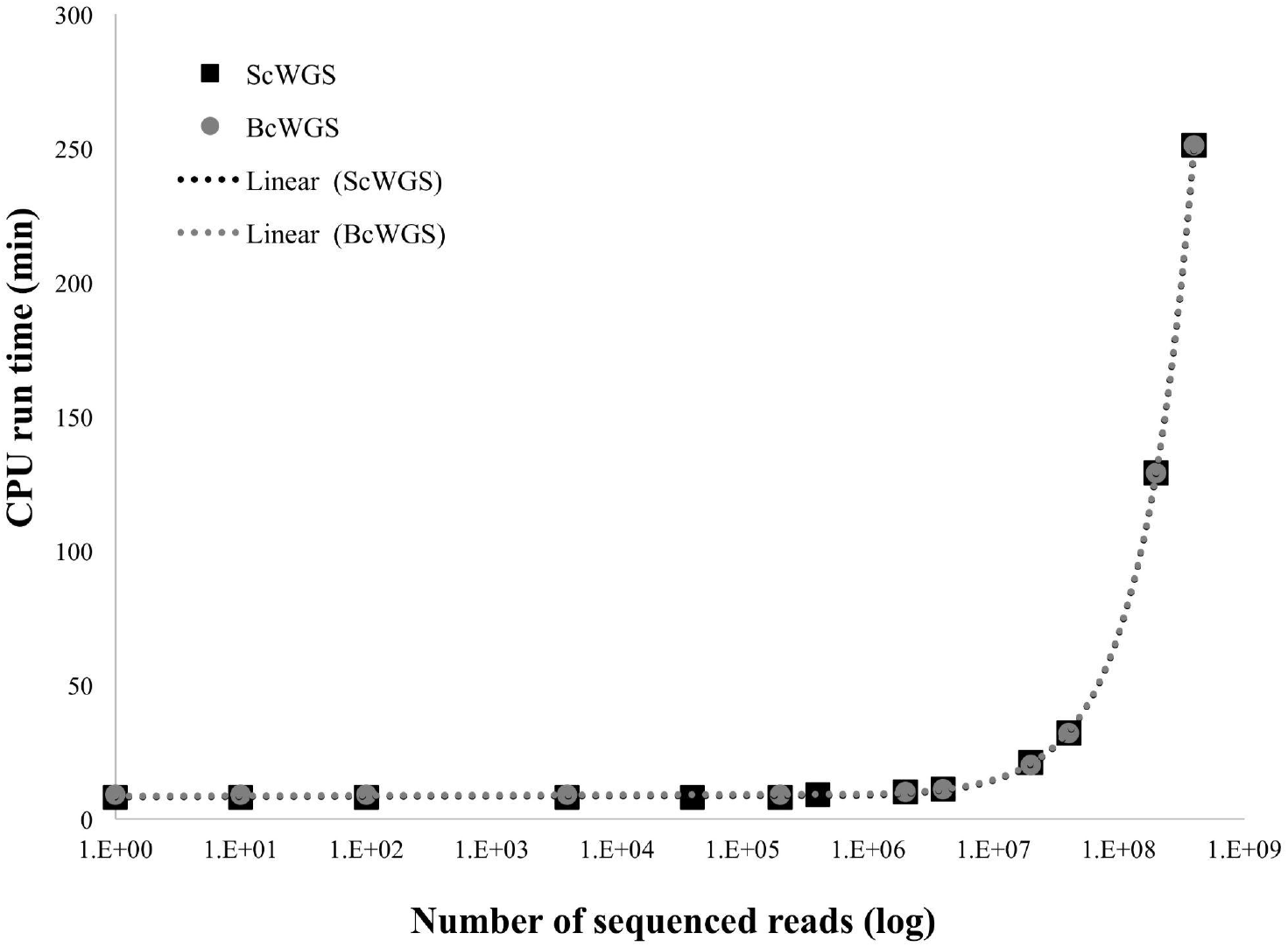
The CPU run time RadiSeq needed to generate different numbers of read sequences in ScWGS and BcWGS simulations are plotted separately. Linear fits to the data are also shown with dotted lines to give a sense of the time requirement for a specific simulation.

Table 1 compares a subset of read alignment statistics, most indicative of the overall trends in data, obtained using Samtools for the RadiSeq simulated data and the experimental data RadiSeq aimed to mimic, for both BcWGS and ScWGS.

**Table 1:**
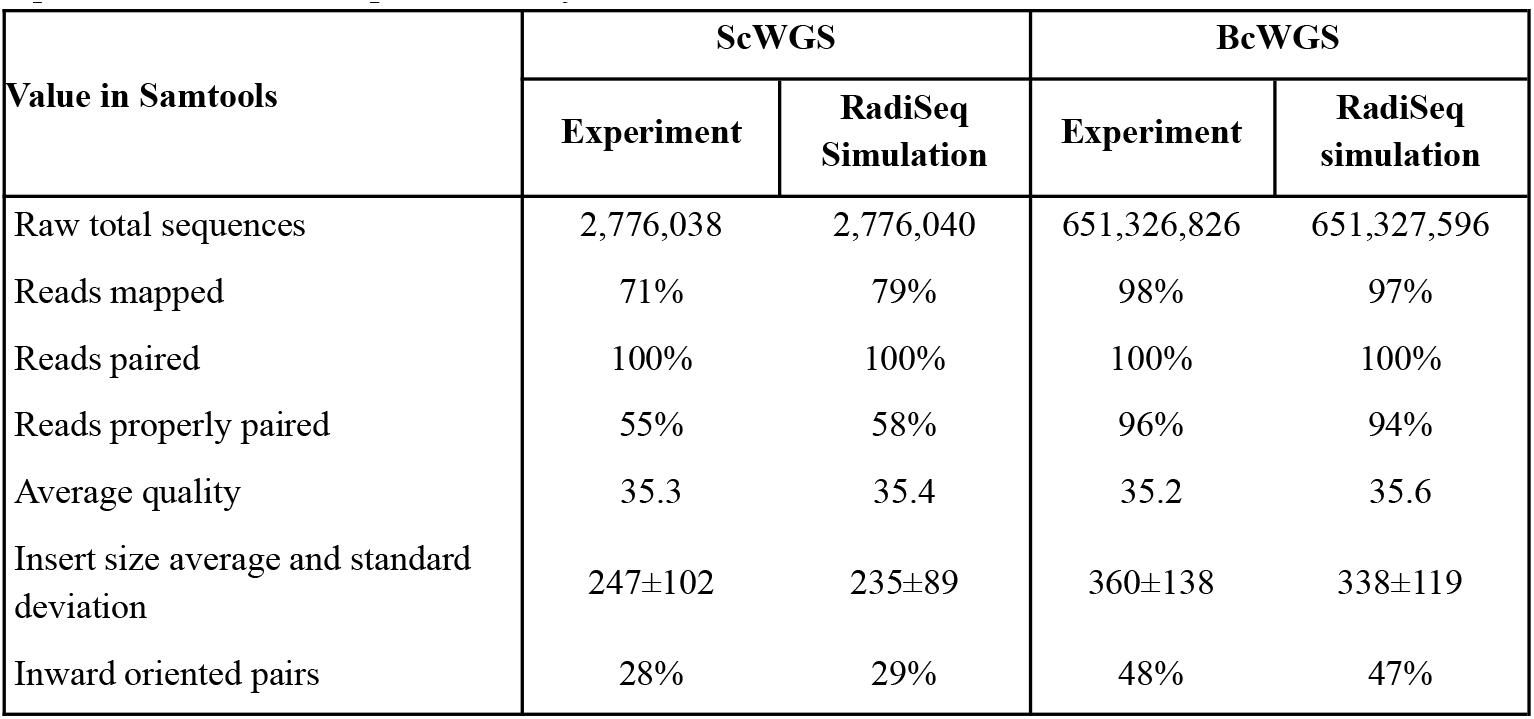
Read alignment statistics obtained using Samtools are compared between the experiment and RadiSeq simulation for both ScWGS and BcWGS.

| Value in Samtools | ScWGS |  | BcWGS |  |
| --- | --- | --- | --- | --- |
|  | Experiment | RadiSeq Simulation | Experiment | RadiSeq simulation |
| Raw total sequences | 2,776,038 | 2,776,040 | 651,326,826 | 651,327,596 |
| Reads mapped | 71% | 79% | 98% | 97% |
| Reads paired | 100% | 100% | 100% | 100% |
| Reads properly paired | 55% | 58% | 96% | 94% |
| Average quality | 35.3 | 35.4 | 35.2 | 35.6 |
| Insert size average and standard deviation | 247±102 | 235±89 | 360±138 | 338±119 |
| Inward oriented pairs | 28% | 29% | 48% | 47% |

In addition to the read-alignment statistics, Samtools provides several underlying distributions that describe the read-aligned data in more detail. In Figure 4, two such distributions: (i) read coverage distribution, and (ii) insert size distribution, are plotted for both the experimentally obtained data and RadiSeq simulated data, for ScWGS and BcWGS separately. A variety of other metrics were also compared (see Supplementary Material) to evaluate the agreement between the simulation and the experiments.

**Figure 4:**
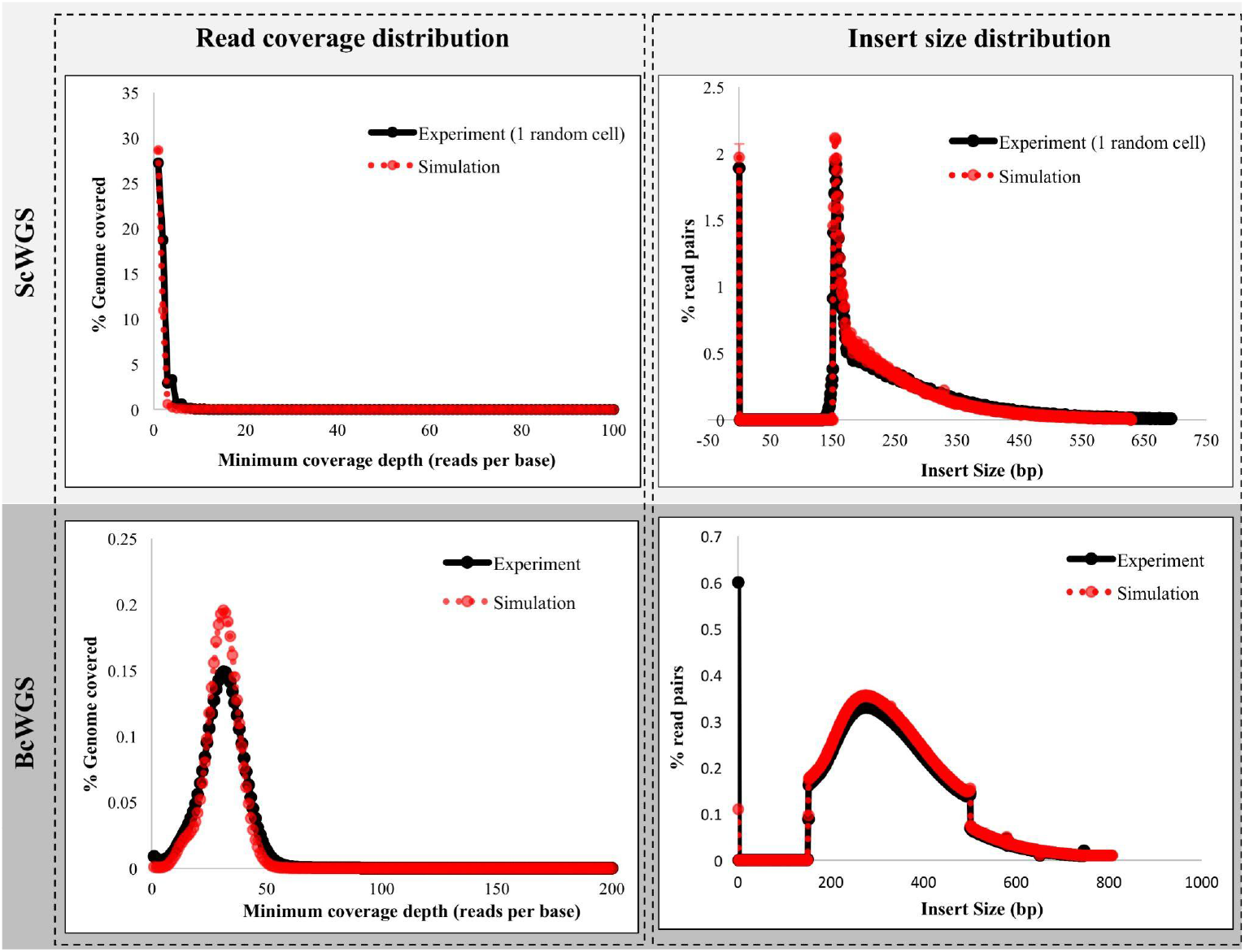
Read coverage distributions (left column) and insert size distributions (right column) obtained from RadiSeq simulation (in red) are compared against the experiment data for single-cell whole-genome sequencing (top row) and bulk-cell whole-genome sequencing (bottom row) separately.

## Discussion

We have built a MC simulation framework called RadiSeq for BcWGS and ScWGS simulation of diploid IR-damaged genomes. To the best of our knowledge, RadiSeq is the first MC framework to simulate the sequencing of radiation-damaged cell models. It is open-source and customizable, and it can generate realistic Illumina sequencing data that can be used for experiment planning and experimental outcome prediction.

Performance testing revealed that simulating the sequencing of sham-irradiated cells takes less than 10 minutes for up to 10^7^ reads with RadiSeq. Beyond this point, the CPU run time increases linearly with the number of reads for both BcWGS and ScWGS simulations, given the same number of CPU threads. For simulations involving up to 10^7^ reads, RadiSeq consistently requires 10 minutes, primarily due to simulation initialization and input file processing. Beyond this threshold, read generation time becomes a more substantial factor, leading to increased overall simulation duration. It is important to note that CPU run time and memory usage can vary significantly based on model customizations, the number of radiation damages in each cell and the computer resources used.

Both the read alignment statistics and the various distributions obtained using Samtools demonstrate reasonable agreement between RadiSeq simulated data and real experimental results for the experimental conditions tested. This demonstrates that RadiSeq can mimic real experiments and generate data that are comparable to real-world sequencing scenarios. While the agreement between the simulated data reported and experimental data is not perfect, it can be improved by fine-tuning the various simulation model parameters. However, accurately modelling experimental variations inherent to intrinsic biological and biochemical techniques remains a challenge.

## Limitations

Overall, RadiSeq demonstrates reasonable computational performance, and acceptable accuracy in simulating read data, closely resembling experimental data. However, RadiSeq has several important limitations.

First, RadiSeq currently only simulates Illumina sequencing chemistry. Although Illumina technology is the most widely used high-throughput next-generation sequencing technology, incorporating other sequencing technologies into the RadiSeq framework as customizable options is desirable and planned for future development.

Second, RadiSeq employs estimations and simplifications of complex biological processes for various model designs, particularly for the MDA amplification model, GC bias model, and the model for read chimeras. As described in the Supplementary Materials, these models are rather rudimentary in the current release of RadiSeq and can be enhanced to incorporate greater complexities.

Third, there is a trade-off between simulation tuning and CPU run time. RadiSeq parameters can be fine-tuned to better match experimental data in order to improve simulation accuracy, but increased accuracy necessitates a longer completion time.

Fourth, the accuracy of RadiSeq in simulating the sequencing of IR-damaged cells has not yet been validated. Doing so will require a range of experimental irradiation conditions with contrasting sequencing outcomes in which experiment and simulation can be compared. Ongoing work by our research group is investigating the variation of sequencing outcomes as a function of radiation dose and radiation type (e.g. photons versus neutrons). Nevertheless, RadiSeq accepts SDD files containing simulated radiation-damaged DNA data and it is thus ready to predict experimental outcomes for post-irradiation DNA sequencing.

Lastly, we acknowledge that while our simplistic technique of folding a damaged geometric genome onto a reference genome is useful for our purpose, it cannot capture the intricacies of mutagenesis. Indeed, simulation of mutagenesis is beyond the scope of RadiSeq and it should be modelled upstream in the MC irradiation pipeline such that the mutations are already encoded in the SDD file that is used as input to RadiSeq. We note that recent work by the TOPAS-nBio collaboration, in developing the MEDRAS (McMahon and Prise 2021) and DaMaRiS (Warmenhoven et al. 2020) packages that model DNA repair, is an important step towards generating mutations from DNA damage. Future work will incorporate the output of these packages into ongoing validation efforts.

## Conclusion

We have developed and released RadiSeq, a first-of-its-kind user-friendly framework to model the single-cell and bulk-cell whole-genome sequencing of IR-damaged DNA. It provides the radiation biology community with a modular and customizable framework that can be used and built upon to aid in experimental design and outcome prediction for post-irradiation sequencing experiments.

## Acknowledgments

We gratefully appreciate the insights and support of Stephan O’Brien (McGill Space Institute) on the computational aspects of this work. Some of the in-built Illumina sequencer error profiles and the read generation methods used in RadiSeq were adopted from the ART Illumina sequencing tool published open-source by the US National Institute of Health (Huang et al. 2012). We sincerely thank the authors Weichun Huang et al. for making the ART toolkit available under an open-source license. This research was enabled in part by computational resources provided by Calcul Quebec (calculquebec.ca) and the Digital Research Alliance of Canada (alliancecan.ca).

## Data availability statement

The source code and user manual for RadiSeq are released open-source under the GPL-3.0 license at https://github.com/kildealab/RadiSeq. Supplementary materials, including detailed methods and results, are accessible online.

## References

Adewoye, Adeolu B., Sarah J. Lindsay, Yuri E. Dubrova, and Matthew E. Hurles. 2015. “The Genome-Wide Effects of Ionizing Radiation on Mutation Induction in the Mammalian Germline.” Nature Communications 6 (March):6684.

Agostinelli, S., J. Allison, K. Amako, J. Apostolakis, H. Araujo, P. Arce, M. Asai, et al. 2003. “Geant4—a Simulation Toolkit.” Nuclear Instruments & Methods in Physics Research. Section A, Accelerators, Spectrometers, Detectors and Associated Equipment 506 (3):250–303.

Alosaimi, Shatha, Armand Bandiang, Noelle van Biljon, Denis Awany, Prisca K. Thami, Milaine S. S. Tchamga, Anmol Kiran, et al. 2020. “A Broad Survey of DNA Sequence Data Simulation Tools.” Briefings in Functional Genomics 19 (1):49–59.

Behjati, Sam, Gunes Gundem, David C. Wedge, Nicola D. Roberts, Patrick S. Tarpey, Susanna L. Cooke, Peter Van Loo, et al. 2016. “Mutational Signatures of Ionizing Radiation in Second Malignancies.” Nature Communications 7 (September):12605.

Bernal, M. A., and J. A. Liendo. 2009. “An Investigation on the Capabilities of the PENELOPE MC Code in Nanodosimetry.” Medical Physics 36 (2):620–25.

Bernal, M. A., D. Sikansi, F. Cavalcante, S. Incerti, C. Champion, V. Ivanchenko, and Z. Francis. 2013. “An Atomistic Geometrical Model of the B-DNA Configuration for DNA–radiation Interaction Simulations.” Computer Physics Communications 184 (12):2840–47.

Bertolet, Alejandro, José Ramos-Méndez, Aimee McNamara, Dohyeon Yoo, Samuel Ingram, Nicholas Henthorn, John-William Warmenhoven, et al. 2022. “Impact of DNA Geometry and Scoring on Monte Carlo Track-Structure Simulations of Initial Radiation-Induced Damage.” Radiation Research 198 (3):207–20.

Chandra, Rohit. 2001. Parallel Programming in OpenMP. Morgan Kaufmann.

Chen, Yen-Chun, Tsunglin Liu, Chun-Hui Yu, Tzen-Yuh Chiang, and Chi-Chuan Hwang. 2013. “Effects of GC Bias in next-Generation-Sequencing Data on de Novo Genome Assembly.” PloS One 8 (4):e62856.

Escalona, Merly, Sara Rocha, and David Posada. 2016. “A Comparison of Tools for the Simulation of Genomic next-Generation Sequencing Data.” Nature Reviews. Genetics 17 (8):459–69.

Faddegon, Bruce, José Ramos-Méndez, Jan Schuemann, Aimee McNamara, Jungwook Shin, Joseph Perl, and Harald Paganetti. 2020. “The TOPAS Tool for Particle Simulation, a Monte Carlo Simulation Tool for Physics, Biology and Clinical Research.” Physica Medica: PM: An International Journal Devoted to the Applications of Physics to Medicine and Biology: Official Journal of the Italian Association of Biomedical Physics 72 (April):114–21.

Ferrari, A., A. Fasso, P. R. Sala, and J. Ranft. 2005. FLUKA: A Multi-Particle Transport Code.

Friedland, Werner, Michael Dingfelder, Pavel Kundrát, and Peter Jacob. 2011. “Track Structures, DNA Targets and Radiation Effects in the Biophysical Monte Carlo Simulation Code PARTRAC.” Mutation Research 711 (1-2): 28–40.

Gawad, Charles, Winston Koh, and Stephen R. Quake. 2016. “Single-Cell Genome Sequencing: Current State of the Science.” Nature Reviews. Genetics 17 (3):175–88.

Huang, Weichun, Leping Li, Jason R. Myers, and Gabor T. Marth. 2012. “ART: A next-Generation Sequencing Read Simulator.” Bioinformatics 28 (4):593–94.

Incerti, S., G. Baldacchino, M. Bernal, R. Capra, C. Champion, Z. Francis, P. Guèye, et al. 2010. “The geant4-DNA Project.” Advances in Complex Systems. A Multidisciplinary Journal 01 (02):157–78.

Kageyama, Shun-Ichiro, Junyan Du, Syuzo Kaneko, Ryuji Hamamoto, Shigeo Yamaguchi, Riu Yamashita, Masayuki Okumura, et al. 2021. “Identification of the Mutation Signature of the Cancer Genome Caused by Irradiation.” Radiotherapy and Oncology: Journal of the European Society for Therapeutic Radiology and Oncology 155 (February):10–16.

Langmead, Ben, and Steven L. Salzberg. 2012. “Fast Gapped-Read Alignment with Bowtie 2.” Nature Methods 9 (4):357–59.

Lasken, Roger S. 2009. “Genomic DNA Amplification by the Multiple Displacement Amplification (MDA) Method.” Biochemical Society Transactions 37 (Pt 2): 450–53.

Li, Heng, Bob Handsaker, Alec Wysoker, Tim Fennell, Jue Ruan, Nils Homer, Gabor Marth, Goncalo Abecasis, Richard Durbin, and 1000 Genome Project Data Processing Subgroup. 2009. “The Sequence Alignment/Map Format and SAMtools.” Bioinformatics 25 (16):2078–79.

Lipman, D. J., and W. R. Pearson. 1985. “Rapid and Sensitive Protein Similarity Searches.” Science 227 (4693):1435–41.

Lu, Na, Yi Qiao, Zuhong Lu, and Jing Tu. 2023. “Chimera: The Spoiler in Multiple Displacement Amplification.” Computational and Structural Biotechnology Journal 21 (February):1688–96.

Mathew, Felix, and John Kildea. 2024. kildealab/RadiSeq: RadiSeq_v2.0. Zenodo. 10.5281/ZENODO.13737532.

Mathew, Felix, James Manalad, Jonathan Yeo, Luc Galarneau, Norma Ybarra, Yu Chang Wang, Patricia N. Tonin, Ioannis Ragoussis, and John Kildea. 2023. “Single-Cell DNA Sequencing-a Potential Dosimetric Tool.” Radiation Protection Dosimetry 199 (15-16): 2047–52.

McMahon, Stephen Joseph, and Kevin M. Prise. 2021. “A Mechanistic DNA Repair and Survival Model (Medras): Applications to Intrinsic Radiosensitivity, Relative Biological Effectiveness and Dose-Rate.” Frontiers in Oncology 11 (June):689112.

McNamara, Aimee L., José Ramos-Méndez, Joseph Perl, Kathryn Held, Naoki Dominguez, Eduardo Moreno, Nicholas T. Henthorn, et al. 2018. “Geometrical Structures for Radiation Biology Research as Implemented in the TOPAS-nBio Toolkit.” Physics in Medicine and Biology 63 (17):175018.

Milhaven, Mark, and Susanne P. Pfeifer. 2023. “Performance Evaluation of Six Popular Short-Read Simulators.” Heredity 130 (2):55–63.

Montgomery, Logan, Christopher M. Lund, Anthony Landry, and John Kildea. 2021. “Towards the Characterization of Neutron Carcinogenesis through Direct Action Simulations of Clustered DNA Damage.” Physics in Medicine and Biology 66 (20). 10.1088/1361-6560/ac2998.

Montgomery, Logan, and James Manalad. 2022. McGillMedPhys/topas_clustered_dna_damage: Indirect Damage Scoring Update. Zenodo. 10.5281/ZENODO.6972469.

Murray, Vincent, Megan E. Hardie, and Shweta D. Gautam. 2019. “Comparison of Different Methods to Determine the DNA Sequence Preference of Ionising Radiation-Induced DNA Damage.” Genes 11 (1). 10.3390/genes11010008.

National Research Council of Canada. Metrology Research Centre. Ionizing Radiation Standards. 2021. EGSnrc: Software for Monte Carlo Simulation of Ionizing Radiation. National Research Council of Canada. 10.4224/40001303.

Nguyen, Van, Irina V. Panyutin, Igor G. Panyutin, and Ronald D. Neumann. 2016. “A Genomic Study of DNA Alteration Events Caused by Ionizing Radiation in Human Embryonic Stem Cells via Next-Generation Sequencing.” Stem Cells International 2016:1346521.

Niita, Koji, Tatsuhiko Sato, Hiroshi Iwase, Hiroyuki Nose, Hiroshi Nakashima, and Lembit Sihver. 2006. “PHITS—a Particle and Heavy Ion Transport Code System.” Radiation Measurements 41 (9-10): 1080–90.

Sakata, Dousatsu, Oleg Belov, Marie-Claude Bordage, Dimitris Emfietzoglou, Susanna Guatelli, Taku Inaniwa, Vladimir Ivanchenko, et al. 2020. “Fully Integrated Monte Carlo Simulation for Evaluating Radiation Induced DNA Damage and Subsequent Repair Using Geant4-DNA.” Scientific Reports 10 (1):20788.

Schuemann, J., A. L. McNamara, J. Ramos-Méndez, J. Perl, K. D. Held, H. Paganetti, S. Incerti, and B. Faddegon. 2019. “TOPAS-nBio: An Extension to the TOPAS Simulation Toolkit for Cellular and Sub-Cellular Radiobiology.” Radiation Research 191 (2):125–38.

Schuemann, J., A. L. McNamara, J. W. Warmenhoven, N. T. Henthorn, K. J. Kirkby, M. J. Merchant, S. Ingram, et al. 2019. “A New Standard DNA Damage (SDD) Data Format.” Radiation Research 191 (1):76–92.

Warmenhoven, John W., Nicholas T. Henthorn, Samuel P. Ingram, Amy L. Chadwick, Marios Sotiropoulos, Nickolay Korabel, Sergei Fedotov, Ranald I. Mackay, Karen J. Kirkby, and Michael J. Merchant. 2020. “Insights into the Non-Homologous End Joining Pathway and Double Strand Break End Mobility Provided by Mechanistic in Silico Modelling.” DNA Repair 85 (January):102743.

Youk, Jeonghwan, Hyun Woo Kwon, Joonoh Lim, Eunji Kim, Taewoo Kim, Ryul Kim, Seongyeol Park, et al. 2024. “Quantitative and Qualitative Mutational Impact of Ionizing Radiation on Normal Cells.” Cell Genomics 4 (2):100499.

Zhao, Min, Di Liu, and Hong Qu. 2017. “Systematic Review of next-Generation Sequencing Simulators: Computational Tools, Features and Perspectives.” Briefings in Functional Genomics 16 (3):121–28.

