## Supplementary file for "RadiSeq: a single- and bulk-cell whole-genome DNA sequencing simulator for radiation-damaged cell models"

Supplementary Material

F. Mathew, Luc Galarneau and J. Kildea

### 1. Introduction

This document contains the supplementary information for the original manuscript "*RadiSeq: a single- and bulk-cell whole-genome sequencing simulator for radiation-damaged cell models*". RadiSeq is an open-source Monte Carlo (MC) simulation framework to perform simulations of whole-genome sequencing of radiation-damaged cells. In this supplementary material, the following aspects are described in more detail: (1) installation instructions of RadiSeq, (2) the sequencing simulation procedure of RadiSeq, (3) the simulation tuning models included in RadiSeq, and (4) the characterization of RadiSeq. This information pertains to the current public release of RadiSeq (RadiSeq_v2.0).

### 2. Installation of RadiSeq

The latest open-source public release of RadiSeq can be downloaded from GitHub (Mathew and Kildea 2024). The current release of RadiSeq requires C++ 17 or above, 64-bit (x64) CPU architecture, and is currently only compatible with Unix-based systems. To get started with RadiSeq, follow the steps below in order.

1. Download the latest release of RadiSeq from GitHub.
2. Unzip the downloaded RadiSeq package.
3. Download a human reference genome of choice.
   RadiSeq will use this reference genome to reconstruct damaged cells during simulations. If unsure, we suggest using the human reference assembly GRCh38/hg38 (Schneider *et al.* 2017) as the default (click here to download)
4. Unzip the downloaded reference genome FASTA file and save it in the 'radiSeqData' folder under the name 'Human_reference_genome.fa'.
5. Compile RadiSeq using the commands:
   - cd *path/to/RadiSeq*
   - make
6. Set up the environment variable 'RADISEQ_DATA_DIR'
   - export RADISEQ_DATA_DIR=*path/to/RadiSeq*/radiSeqData

   You will need to do this step every time you open a fresh Terminal window. Alternatively, you may choose to add this to one of your startup files (e.g. *.bashrc*) if you are comfortable doing so.

Note: Replace '*path/to/RadiSeq*' in steps 5 and 6 with the actual full path to the directory RadiSeq in your system

More instructions and details on installation, running a test simulation, and using the Illumina sequencer profiler tool can be found on GitHub.

### 3. Simulation Overview

This section provides an overview of the RadiSeq simulation framework. It is organized into three subsections: (i) Setting up a Simulation in RadiSeq, describing the input files and expected formatting of these files (ii) RadiSeq Simulation Procedure, which outlines the step-by-step methodology employed by RadiSeq to generate simulated sequencing data, and (iii) RadiSeq Pipeline Schematic, which presents a visual representation of the workflow and components within the RadiSeq pipeline.

#### 3.1. Setting up a Simulation in RadiSeq

Setting up a sequencing simulation is straightforward once RadiSeq is successfully installed and tested. To run a simulation, users simply need to specify the path to a parameter text (.txt) file as the only argument. This file contains all the necessary input parameters for configuring the simulation, including the paths to the other input files that are needed.

##### 3.1.1. User Parameter File

Users can create a parameter file with any filename, provided that the contents are formatted as specified name-value pairs. Input values should be given in the following format: Parameter Name = Parameter Value #comment. Table 1 lists all acceptable parameter names, and Figure 1 shows a screenshot of an example parameter file. All parameters are optional except for the path to the Standard DNA Damage (SDD) file (Schuemann et al. 2019). Parameters other than the path to the SDD file have default values, as shown in Table 1. However, any user-provided value will override the defaults during a simulation run.

Certain parameters remain inactive until their dependent parameters are activated. For example, the parameters specifying the number of SDD files to combine damages from ("*number_of_particles_to_merge*"), the name of particles used in the irradiation simulation to generate the SDD files ("*primary_particles_simulated*") will remain inactive until the parameter determining whether damages from multiple particles need to be combined ("*merge_damages_from_multiple_particles*") is set to 'True'. RadiSeq will disregard inactive parameter values and enforce proper settings for active parameters when needed during a simulation run.

***Table 1****: List of all adjustable parameters, their descriptions and acceptable value types in RadiSeq. Default parameter values are indicated in bold font unless explicitly specified in the Description column. Parameters that do not have default values as their dependent parameters are deactivated by default and are indicated with a *.*

| **Parameter name** | **Description** | **Acceptable value types** |
|---|---|---|
| sddFilePath | The full path to the SDD file(s). Mandatory parameter. | Comma-separated list of file paths (string) |
| merge_damages_from_multiple_particles | Flag to indicate if the user wishes to define a single genome by combining damages from multiple SDD files | 'True' or '**False**' |

| | | |
|---|---|---|
| number_of_particles_to_merge | Number of SDD files from individual primary simulations to combine | *Number (integer) |
| primary_particles_simulated | Names of primary particles that introduced the damages that are going to be combined into a single genome | *Comma-separated list of names (string) |
| adjust_damages_with_actual_dose | Flag to indicate if the user wishes to scale the number of damages with the actual dose delivered and it is different from the expected dose | 'True' or '**False**' |
| actual_dose_delivered_data | The full path to the file (.txt) containing the actual dose delivered in each exposure. One file is expected for each SDD file specified | *Comma-separated list of paths (string) |
| reference_genome_FASTAfile | The full path to the reference genome file. Defaults to the genome included during installation. Genomes of any diploid organism can be specified. | Path to file (string) |
| acceptable_difference_in_seq_length_percent | Acceptable difference in the lengths of the reference genome provided and the genome length of the Monte Carlo model. Default is 10%. | Percentage (double) |
| number_of_cells_in_sample | Total number of cells you assume to have in your sample flask. This is different from the number of cells to sequence. Default is 10. | Number (integer) |
| number_of_cells_to_sequence | Number of cells (damaged and undamaged) to be sequenced. These many cells will be randomly selected from the number of cells in the sample. Default is 5. | Number (integer) |
| illumina_sequencer | Name of the Illumina sequencer to be used for sequencing from the in-built list | 'HiSeq1000', 'HiSeq2000', 'HiSeq2500_v125', 'HiSeq2500_v150', '**HiSeqX**', 'NovaSeq6000' and |

| | | |
|---|---|---|
| | | 'Custom' |
| custom_read1_quality_profile_path | The full path to the read 1 quality profile file when the custom sequencer is chosen | *Path to file (string) |
| custom_read2_quality_profile_path | The full path to the read 2 quality profile file when the custom sequencer is chosen and paired-end sequencing is needed | *Path to file (string) |
| single_or_bulk_sequencing | Flag to specify if single-cell or bulk-cell sequencing is to be performed | '**single**' or 'bulk' |
| do_paired_end_sequencing | Flag to indicate if paired-end sequencing is to be performed | '**True**' or 'False' |
| fraction_of_other_oriented_read_pairs | The fraction of read pairs needs to be in orientations other than forward-reverse (FR). Default is 0. | Number in the range [0,1] (double) |
| fragment_size_distribution_path | The full path to the text (.txt) file that stores the fragment size distribution | *Path to file (string) |
| min_DNA_fragment_length | Minimum DNA fragment length (in bp) to be generated if paired-end sequencing. Default is 200. | Number (integer) |
| max_DNA_fragment_length | Maximum DNA fragment length (in bp) to be generated if paired-end sequencing. Default is 800. | Number (integer) |
| mode_DNA_fragment_length | Mode DNA fragment length (in bp) to be generated if paired-end sequencing. Default is 400. | Number (integer) |
| beta_of_beta_distribution | The beta parameter value for the beta distribution that will be used to represent the fragment size distribution. Default is 4.5. | Number (integer) |
| read_length | Length of the read (in bp) to be generated | *Number (integer) |
| total_read_coverage | Total read coverage the user wants to get from this sequencing. If | Number (integer) |

| | | |
|---|---|---|
| | single-cell sequencing the read coverage will get distributed over the total number of cells sequenced. Default is 0.01. | |
| coverage_distribution | Read-coverage distribution mode to be used in single-cell sequencing | '**Uniform**' or 'MDA' |
| degree_of_GC_bias | The slope of the linear portions of the triangular function used for GC bias. Default is 0. | Number (double) |
| bin_size_for_GC_bias_estimation | The bin size to be used to calculate the GC fraction and bias. Default is 10,000. | Number (integer) |
| read1_insertion_error_rate | The insertion error rate for read 1. Default is 0.00002 | Number in the range [0,1] (double) |
| read1_deletion_error_rate | The deletion error rate for read 1. Default is 0.00011 | Number in the range [0,1] (double) |
| read2_insertion_error_rate | The insertion error rate for read 2. Default is 0.00009 | Number in the range [0,1] (double) |
| read2_deletion_error_rate | The deletion error rate for read 2. Default is 0.00023 | Number in the range [0,1] (double) |
| read_artifacts_rate | The rate of chimera artifact formation in read 1 and 2 combined. Default is 0. | Number in the range [0,1] (double) |
| output_directory_path | The full path to the directory where the output FASTQ files and the run summary file should be stored. Default is './output' | Path to directory (string) |
| output_FASTQ_filename_prefix | Prefix for the sequenced output FASTQ file (omit file extension). Default is 'CELL' | String |
| make_summary_report | Flag to indicate if the user wishes to generate a summary report file at the end of a run | '**True**' or 'False' |
| random_seed | Seed for the random number generator to be initialized with a fixed seed. A default value of 0 | Number (integer) |

| | | |
|---|---|---|
| | indicates that the system will automatically generate seeds randomly | |
| number_of_threads | Number of threads to be used for a multithreaded run. The default value is 1 | Number (integer) |

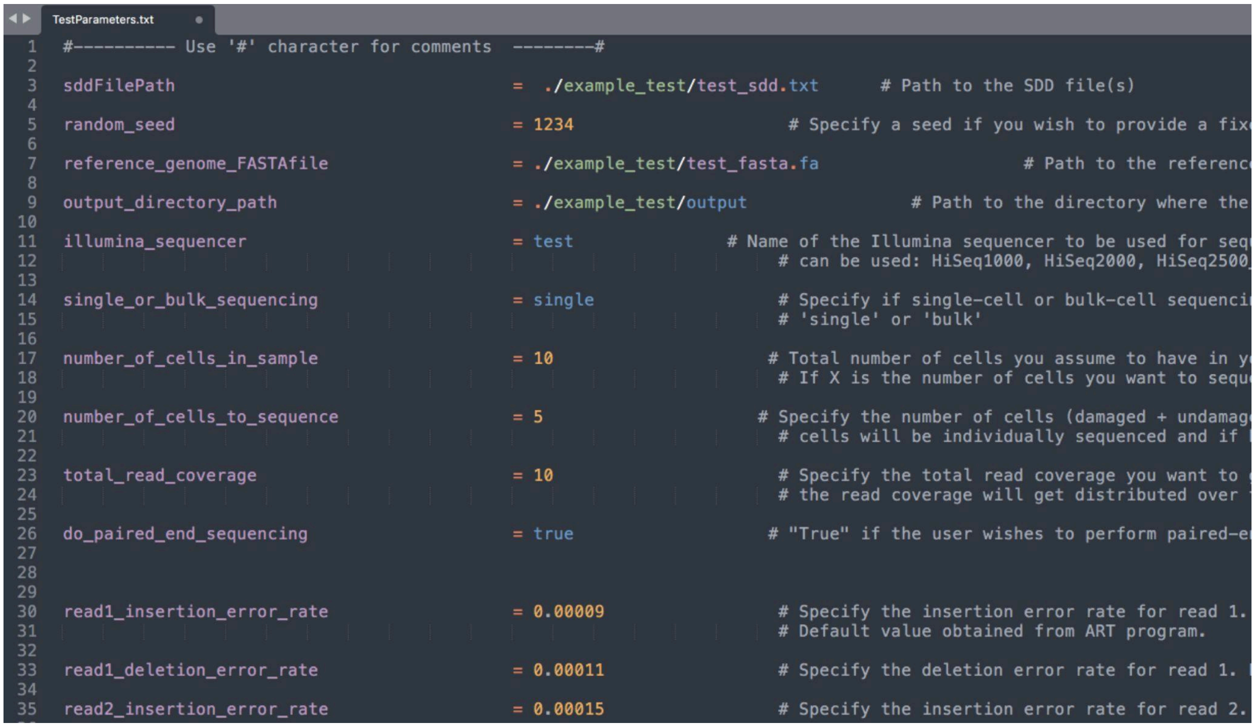

```
TestParameters.txt
#---------- Use '#' character for comments  --------#

sddFilePath                         =  ./example_test/test_sdd.txt      # Path to the SDD file(s)

random_seed                         = 1234             # Specify a seed if you wish to provide a fix

reference_genome_FASTAfile          = ./example_test/test_fasta.fa                # Path to the referenc

output_directory_path               = ./example_test/output               # Path to the directory where the

illumina_sequencer                  = test             # Name of the Illumina sequencer to be used for seq
                                                            # can be used: HiSeq1000, HiSeq2000, HiSeq2500_

single_or_bulk_sequencing           = single                # Specify if single-cell or bulk-cell sequenci
                                                            # 'single' or 'bulk'

number_of_cells_in_sample           = 10                   # Total number of cells you assume to have in y
                                                            # If X is the number of cells you want to sequ

number_of_cells_to_sequence         = 5                   # Specify the number of cells (damaged + undamag
                                                            # cells will be individually sequenced and if

total_read_coverage                 = 10                    # Specify the total read coverage you want to
                                                            # the read coverage will get distributed over

do_paired_end_sequencing            = true                 # "True" if the user wishes to perform paired-e

read1_insertion_error_rate          = 0.00009               # Specify the insertion error rate for read 1.
                                                            # Default value obtained from ART program.

read1_deletion_error_rate           = 0.00011               # Specify the deletion error rate for read 1.

read2_insertion_error_rate          = 0.00015               # Specify the insertion error rate for read 2.
```

***Figure 1:*** *Screenshot showing the contents of a sample parameter file for RadiSeq. This sample shows some of the different parameters a user can set in a key-value format.*

Except for the SDD file and the reference genome FASTA file, which have established formatting standards, all other user input files for RadiSeq must follow specific formatting guidelines to ensure the simulation is set up correctly. Users are responsible for verifying that these files adhere to the formatting specifications outlined here.

3.1.2. Actual Dose-Delivered Data File

When the parameter to scale the number of damages in an SDD file based on the actual dose delivered during the irradiation simulation ("*adjust_damages_with_actual_dose*") is set to 'True', RadiSeq requires users to provide the actual dose-delivered data file using the input parameter "*actual_dose_delivered_data*". For each SDD file, a corresponding actual dose-delivered data file is expected. These files must be in simple text format (.txt), and their content should contain the numerical value of the actual dose delivered during irradiation, measured in Gray (Gy). If the

SDD file contains multiple exposure (irradiation) records, the dose-delivered data file must have the same number of rows as exposures in the SDD file, with each row specifying the dose-delivered for the corresponding exposure.

##### 3.1.3. Custom Read Quality Profile File

Custom read quality profile files are required for both read 1 and read 2 when a 'custom' sequencer is selected as the Illumina sequencer for the simulation using the parameter "*illumina_sequencer*". These files can be specified using the parameters "*custom_read1_quality_profile_path*" and "*custom_read2_quality_profile_path*", respectively. RadiSeq will only accept read-quality profiles in a specific format. Users can generate these files using the RadiSeqProfiler tool, which is included in the RadiSeq package. Instructions on how to use this tool to generate quality profile files for custom sequencers can be found on GitHub.

##### 3.1.4. Fragment Size Distribution Data File

The DNA fragment size distribution file can be specified using the parameter "*fragment_size_distribution_path*". This file must be a simple text file (.txt) with multiple rows of data entries. Each row should contain two numbers, arranged as a key-value pair, separated by a tab. The first number represents the length of the DNA fragment, and the second indicates the fraction of DNA fragments with that specific length. The order of the key-value pairs in the file can be random. If the user provides discontinuous DNA fragment lengths, RadiSeq will perform linear interpolation to estimate the missing data pairs and renormalize the DNA fragment fractions to sum to 1.

#### 3.2. RadiSeq Simulation Procedure

The first task in RadiSeq is processing the input parameters from the parameter file provided as an argument. RadiSeq verifies that all required parameters are included and checks whether appropriate values are assigned. If a required parameter is missing, RadiSeq will terminate with an error message indicating the missing parameter. If an optional parameter is not provided, RadiSeq will proceed using default values. Providing the appropriate parameters is the only way for users to configure the simulation according to their preferences.

The sequencing simulation of a radiation-damaged cell in RadiSeq begins with the reconstruction of its genome based on the geometric damage information in the SDD file. Any MC geometric nuclear DNA model compatible with recording radiation-induced DNA damage in the SDD file format can be used for this process. For our simulations, we used the in-house developed NICE geometric nuclear DNA model (Montgomery *et al.* 2021), built in TOPAS-nBio, to perform cell irradiation simulations and generate SDD files of damaged cells. SDD files are designed to store damage records from multiple repeated irradiation simulations (exposures), provided the irradiation conditions remain unchanged. These repeated simulation records can represent multiple cells in the sample, each with its own unique DNA damage profile.

In MC cell irradiation simulations, exposure from each radiation type is often simulated independently for ease of simulation, although, in reality where secondary particles are encountered, the cell may experience multiple radiation types simultaneously. In such cases, each radiation type generates its own SDD file, making it necessary to combine the geometric damage information from multiple SDD files to obtain a complete description of the damaged genome. To address this, RadiSeq provides the option to specify multiple SDD files, each corresponding to the irradiation of the same geometric DNA model with a different radiation type. Additionally, users can scale down the number of damages linearly with the actual dose exposed if needed during genome reconstruction using the "*adjust_damages_with_actual_dose*" parameter.

When a simulation is initialized with an SDD file, RadiSeq scans the file to determine the number of simulation records (referred to here as damaged cells) it contains. If the number of cells specified by the user (using the parameter "*number_of_cells_in_sample*") exceeds the number of damaged cells available in the SDD file, RadiSeq will assume that the remaining cells are undamaged. Decisions such as this are promptly communicated to the user during runtime and included in the simulation run summary. Before starting the reconstruction of damaged genomes, RadiSeq checks for sufficient memory to temporarily store the reconstructed genomes and proceeds with the simulation only if adequate memory is available.

RadiSeq starts the reconstruction of both damaged and undamaged cells using the reference genome specified in the parameter file ("*reference_genome_FASTAfile*"), applying the damages recorded in the SDD file to the double-stranded genome. If multiple SDD files corresponding to different radiation types are provided, the damages from each file will be combined for each cell in the order in which they are listed in the SDD files. That is, the $n^{th}$ irradiation record in each SDD file is assumed to correspond to the same $n^{th}$ cell. Not all cells in an SDD file may need to be reconstructed; only the number of cells specified for sequencing (using the parameter "*number_of_cells_to_sequence*") will be reconstructed in RadiSeq. During reconstruction, a break is introduced in the genome sequence where a strand break is needed, fragmenting the genome. Additionally, any base damage is recorded by replacing the nitrogen base with an 'N'. The reconstructed genomes are temporarily stored in memory until the end of the simulation run, after which RadiSeq is ready to start the sequencing.

Based on the sequencing procedure chosen (using the parameter "*single_or_bulk_sequencing*"), i.e., bulk-cell whole genome sequencing (BcWGS) or single-cell whole genome sequencing (ScWGS), RadiSeq will access the cell data differently. For ScWGS, each cell that needs to be sequenced will be read one by one, whereas, for BcWGS, cells are randomly accessed from the sample bulk to create a sequenced read.

The first step in generating sequenced read data from a cell is to determine the location for read creation along the genome, known as the initial primer binding site. In RadiSeq, the selection of the initial primer binding site depends on: (i) DNA amplification bias—over-representation of certain genomic regions due to biased amplification, (ii) GC bias—preference for genomic regions rich in or lacking G and C bases, and (iii) the availability of continuous sequence for the desired read length or read pair (i.e., no fragmentation or chromosome discontinuity). Users can choose between available amplification bias models using the parameter "*coverage_distribution*", set the GC bias with "*degree_of_GC_bias*", and specify DNA fragmentation using "*fragment_size_distribution_path*" or the four parameters describing a beta function ("*min_DNA_fragment_length*", "*max_DNA_fragment_length*", "*mode_DNA_fragment_length*", and "*beta_of_beta_distribution*"). For a uniform genome amplification procedure, the initial primer binding site is determined randomly. However, if GC bias is specified, the initial primer binding site selection is influenced by the GC fraction of each genomic segment and the provided bias.

If paired-end sequencing is needed (specified using the "*do_paired_end_sequencing*" parameter), the length of the DNA fragment is determined using the DNA fragmentation model, and the location of the second read in the pair is identified as the end of the fragment. The sequence corresponding to the length of a read is then read according to the read orientation model, which can be tuned using the parameter "*fraction_of_other_oriented_read_pairs*", in appropriate orientation as template sequences.

The final step of read generation is to include sequencing errors and artifacts in the read templates. Insertions and deletions are randomly made in the read according to the rates of insertions and deletions, which can differ for the two reads in a pair. Based on the Illumina (Illumina Inc. San Diego, California, USA) sequencer error model, determined using the chosen sequencer ("*illumina_sequencer*"), RadiSeq will generate a quality profile for the read. At locations in the read where the quality is low, a base substitution is introduced at random. Additionally, the quality values corresponding to bases marked as 'N' are set to a minimum. Finally, chimeras are introduced to the reads according to the chimeric read formation model (see Supplementary Methods), which can be tuned using the "*read_artifacts_rate*" parameter.

The final read information and the corresponding quality profile are written in a FASTQ file and compressed. RadiSeq will generate two fastq.gz files if paired-end sequencing is performed. For ScWGS, each cell will have its own set of fastq.gz files. Additionally, RadiSeq generates a run summary file ("*make_summary_report*") that contains all the descriptions of the simulation performed.

#### 3.3. RadiSeq Pipeline Schematic

Figure 2 shows a detailed schematic of the RadiSeq pipeline. The input parameter file is the only explicit input to RadiSeq, and all other input parameters must be included in this file. The RadiSeq simulation proceeds through four steps: Step 1: Damaged Genome Reconstruction, Step 2: Sequencing Simulation Initiation, Step 3: Sequencing Simulation Tuning, and Step 4: Read Generation. At the end of the simulation, RadiSeq generates two output files.

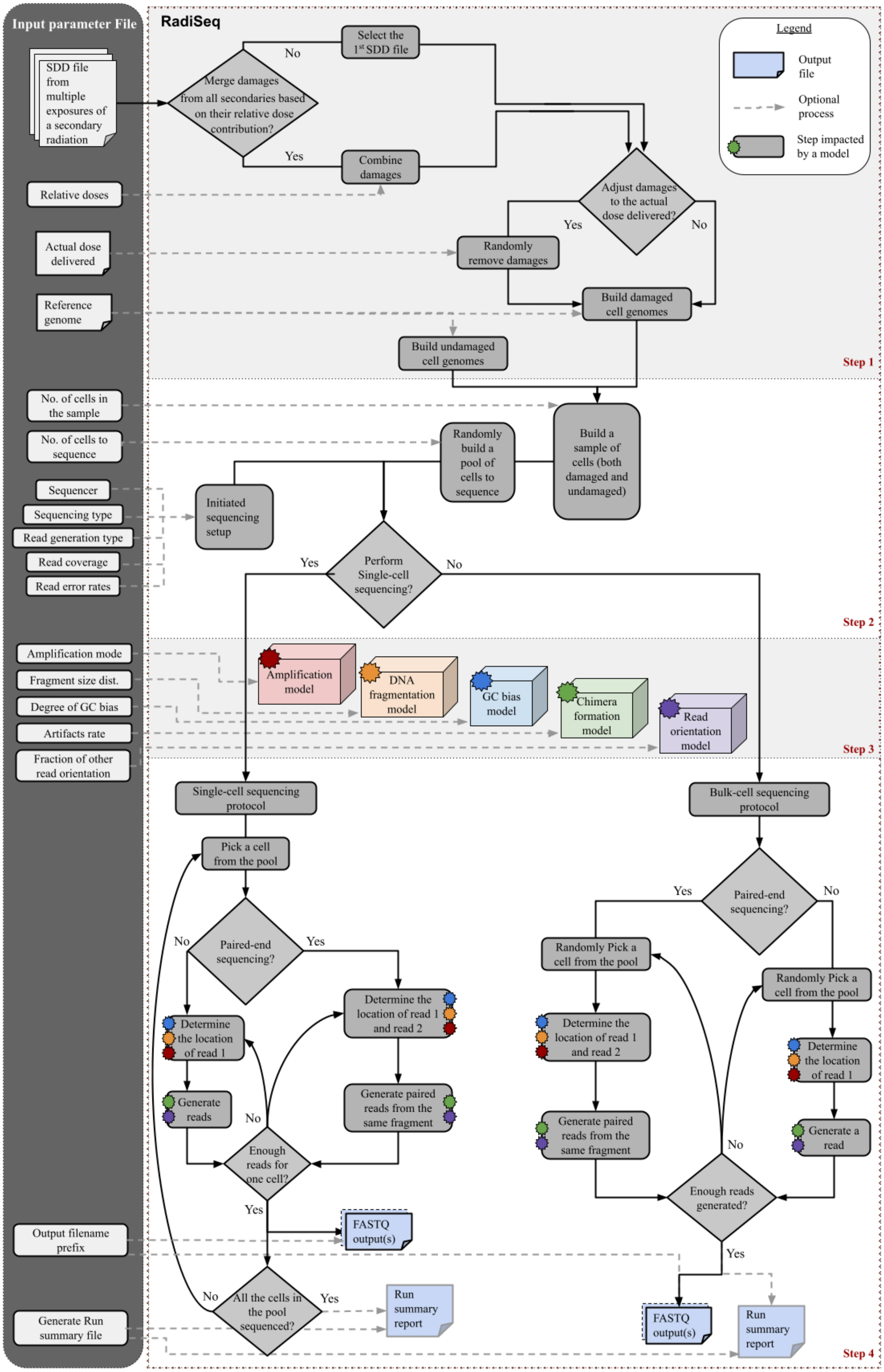

Input parameter File
SDD file from multiple exposures of a secondary radiation
Relative doses
Actual dose delivered
Reference genome
No. of cells in the sample
No. of cells to sequence
Sequencer
Sequencing type
Read generation type
Read coverage
Read error rates
Amplification mode
Fragment size dist.
Degree of GC bias
Artifacts rate
Fraction of other read orientation
Output filename prefix
Generate Run summary file
RadiSeq
Merge damages from all secondaries based on their relative dose contribution?
No
Yes
Select the 1st SDD file
Combine damages
Adjust damages to the actual dose delivered?
Randomly remove damages
Build damaged cell genomes
Build undamaged cell genomes
Legend
Output file
Optional process
Step impacted by a model
Step 1
Build a sample of cells (both damaged and undamaged)
Randomly build a pool of cells to sequence
Initiated sequencing setup
Perform Single-cell sequencing?
Step 2
Amplification model
DNA fragmentation model
GC bias model
Chimera formation model
Read orientation model
Step 3
Single-cell sequencing protocol
Pick a cell from the pool
Paired-end sequencing?
Determine the location of read 1
Generate reads
Determine the location of read 1 and read 2
Generate paired reads from the same fragment
Enough reads for one cell?
FASTQ output(s)
All the cells in the pool sequenced?
Run summary report
Bulk-cell sequencing protocol
Randomly Pick a cell from the pool
Generate a read
Enough reads generated?
Step 4

***Figure 2**: Schematic showing the pipeline of the RadiSeq simulation framework. The four steps in RadiSeq simulation (Step 1: Damaged Genome Reconstruction, Step 2: Sequencing Simulation Initiation, Step 3: Sequencing Simulation Tuning, and Step 4: Read Generation) are indicated.*

### 4. Supplementary Methods

This section provides additional details on RadiSeq simulation aspects not described in the main manuscript, including details on different simulation tuning models.

#### 4.1. Random Number Generator

RadiSeq heavily relies on pseudo-random numbers throughout the sequencing simulation. These numbers are essential for processes such as selecting a random genomic location for read generation, choosing a cell to be sequenced from the sample, deciding the base to be substituted in the genome, and generating a DNA fragment size. RadiSeq employs the Mersenne Twister algorithm (Matsumoto and Nishimura 1998) from the C++ standard library as its Random Number Generator (RNG) and ensures it is thread-safe to maintain compatibility with multithreading. Users have the option to manually set a fixed seed or opt for a random seed for the RNG using the parameter "*random_seed*". The random seed is generated as a seed sequence of eight numbers, which includes a high-resolution clock time and seven random numbers from the random device class of C++.

#### 4.2. Illumina Sequencer Models

RadiSeq simulates the Illumina sequencing procedure for read generation and error insertion, requiring error profiles of the sequencers users wish to emulate. RadiSeq includes error profiles for five Illumina sequencers: HiSeq 1000, HiSeq 2000, HiSeq 2500, HiSeqX, and NovaSeq 6000. Except for the NovaSeq 6000 profile that we generated ourselves, these profiles were adopted from the ART simulation tool available as open-source (Huang *et al.* 2012). Users can choose from the available list of sequencer models using the parameter "*illumina_sequencer*" and that will determine the error profile to be used. RadiSeq also provides the option to include any other Illumina sequencer profile of choice by allowing users to add a custom error profile ("*custom_read1_quality_profile_path*"). Users can generate their own profiles using the RadiSeqProfiler tool included in the toolkit.

#### 4.3. Multiple Displacement Amplification Model

Uniform DNA amplification of the genome can be easily modelled in RadiSeq using its default settings. This uniform amplification is emulated by a random genome sampling approach during read generation in RadiSeq, assuming an equal probability for every region of the genome to be sequenced, unless a bias such as the GC bias model is applied in the simulation. The uniform DNA amplification model can mimic experimental approaches such as the polymerase chain reaction (PCR) (Erlich *et al.* 1989) and primary template-directed amplification (PTA) (Gonzalez-Pena *et al.* 2021) techniques, where unbiased amplification is expected.

RadiSeq also includes a biased amplification model representing the classic multiple displacement amplification (MDA) (Spits *et al.* 2006) approach. MDA is an isothermal amplification procedure suitable for amplifying small amounts of genomic material, such as a single-cell genome. The exponential amplification of MDA is associated with sequence-dependent overamplification, leading to highly non-uniform amplification bias, where random regions of the genome are sequenced more than others. In RadiSeq, this bias is implemented in a two-step process. First, the number and location of initial primer binding sites are determined randomly. Then, a random number of these initial primer sites are identified as overamplification sites and are sequenced more frequently, generating an MDA-like coverage distribution.

One limitation of the current implementation of amplification models in RadiSeq is that they do not account for sequencing error propagation during amplification. In reality, sequencing errors occurring in early amplicons can be carried forward to subsequent amplicons, potentially introducing differences between the sequencing profile of the simulated data and the experiment, especially for MDA. Therefore, caution must be taken when performing variant calling analysis on the simulated data. Error propagation is omitted in the current version of RadiSeq due to the computational expense and time requirements but can be added in the later releases.

#### 4.4. DNA Fragmentation Model

To account for differences in DNA fragmentation between different experiments, we have included two ways to specify the required DNA fragment size distribution in RadiSeq simulations. The first approach uses a Beta distribution (Gupta and Nadarajah 2004), shown in equation 1, to characterize the fragment size distribution. Users can modify the upper and lower bounds (maximum and minimum fragment sizes), the mode value of the fragment size distribution, and the beta value, which together determine the shape of the distribution.

$$B(\alpha, \beta) = \frac{\Gamma(\alpha)\Gamma(\beta)}{\Gamma(\alpha+\beta)} \qquad \text{Eq. 1}$$

Where $B(\alpha, \beta)$ is the beta probability distribution function defined using the $\alpha$ and $\beta$ parameters. $\Gamma$ is the gamma function. The $\alpha$ parameter value is derived from the equation for the mode of the beta distribution, referred to here as betaMode, from the upper and lower bounds, mode value and $\beta$ values using equations 2 and 3.

$$\text{betaMode} = \frac{\alpha-1}{\alpha+\beta-2} \qquad \text{Eq. 2}$$

$$\text{betaMode} = \frac{\textit{mode value} - \textit{lower bound}}{\textit{upper bound} - \textit{lower bound}} \qquad \text{Eq. 3}$$

The second option is more direct: users can specify the path to a file that stores any distribution for RadiSeq to load. The data in this external file must be formatted in key-value pairs separated by a tab in each row, with the first value being the fragment length and the second value the frequency of DNA fragments of that specific length. As previously mentioned, if the user provides discontinuous DNA fragment lengths, RadiSeq will perform linear interpolation to estimate the missing data pairs and renormalize the DNA fragment fractions to sum to 1.

#### 4.5. GC Bias Model

An ideally unbiased genome would have an almost equal contribution of the nitrogen bases adenine (A), thymine (T), guanine (G), and cytosine (C) in its composition. However, in actual genomes, we can observe regions that are AT-rich (with more A and T than G and C) and GC-rich (with more G and C than A and T) (Thakur, Packiaraj and Henikoff 2021). GC bias refers to the positive or negative correlation in sequencing coverage of GC-enriched or GC-deficient regions (Lynch *et al.* 2010). Experiments have shown both positive and negative GC biases, with varying degrees depending on the protocols and sequencing kits used. However, a unimodal bias (Benjamini and Speed 2012) is assumed to be more representative of real scenarios (Chen *et al.* 2013). In RadiSeq, this unimodal bias is constructed as a combination of two linear components using a simple triangular function given in equation 4. The degree of bias can be controlled by adjusting the slope of the linear components (using the parameter "*degree_of_GC_bias*").

$$F(\text{GC fraction}) = \begin{cases} (\text{Degree of GC bias} \times (\text{GC fraction} - 0.5) \times 100) + 1; & \text{GC fraction} <= 0.5 \\ (-\text{Degree of GC bias} \times (\text{GC fraction} - 0.5) \times 100) + 1; & \text{GC fraction} > 0.5 \end{cases} \quad \text{Eq. 4}$$

#### 4.6. Chimeric Read Formation Model

Chimeric sequences (chimeras) are formed by the misjoining of two or more non-adjacent genomic regions. These amplification artifacts occur more frequently in MDA (Lu *et al.* 2023). Chimeras can be classified into two types: (1) inverted chimeras and (2) direct chimeras. The simple model of chimeric read formation in RadiSeq can only generate inverted chimeras. The chimeric model first identifies a region in the DNA fragment as a chimeric and inserts a reverse complementary sequence to that segment to form an inverted chimera. The length of the chimeric region and the location are both randomly determined during the simulation run. However, users can modify the rate of these chimera formations using an input parameter ("*read_artifacts_rate*") to tailor to their simulation needs. Changing the number of chimeras can potentially impact the number of reads mapped successfully and the number of reads properly paired.

#### 4.7. Read Orientation Model

In paired-end sequencing, the two read pairs can have multiple orientations, such as forward-reverse (FR) and reverse-forward (RF) as shown in Figure 3. Different sequencing experiments can exhibit varying proportions of these read pair orientations due to factors like sequencing artifacts and inversion mutations. RadiSeq includes an option to generate different proportions of RF, FR and other (FF and RR) read orientations ("*fraction_of_other_oriented_read_pairs*"), which can influence the number of reads that are properly paired and mapped to the reference genome. RadiSeq controls the read orientation by manipulating how the read templates are read from a DNA fragment. However, this artificial modification of read pair orientation does not result from underlying sequence alterations. Therefore, the simulated data should not be used to interpret results from read orientation-based variant analysis (Diossy *et al.* 2021).

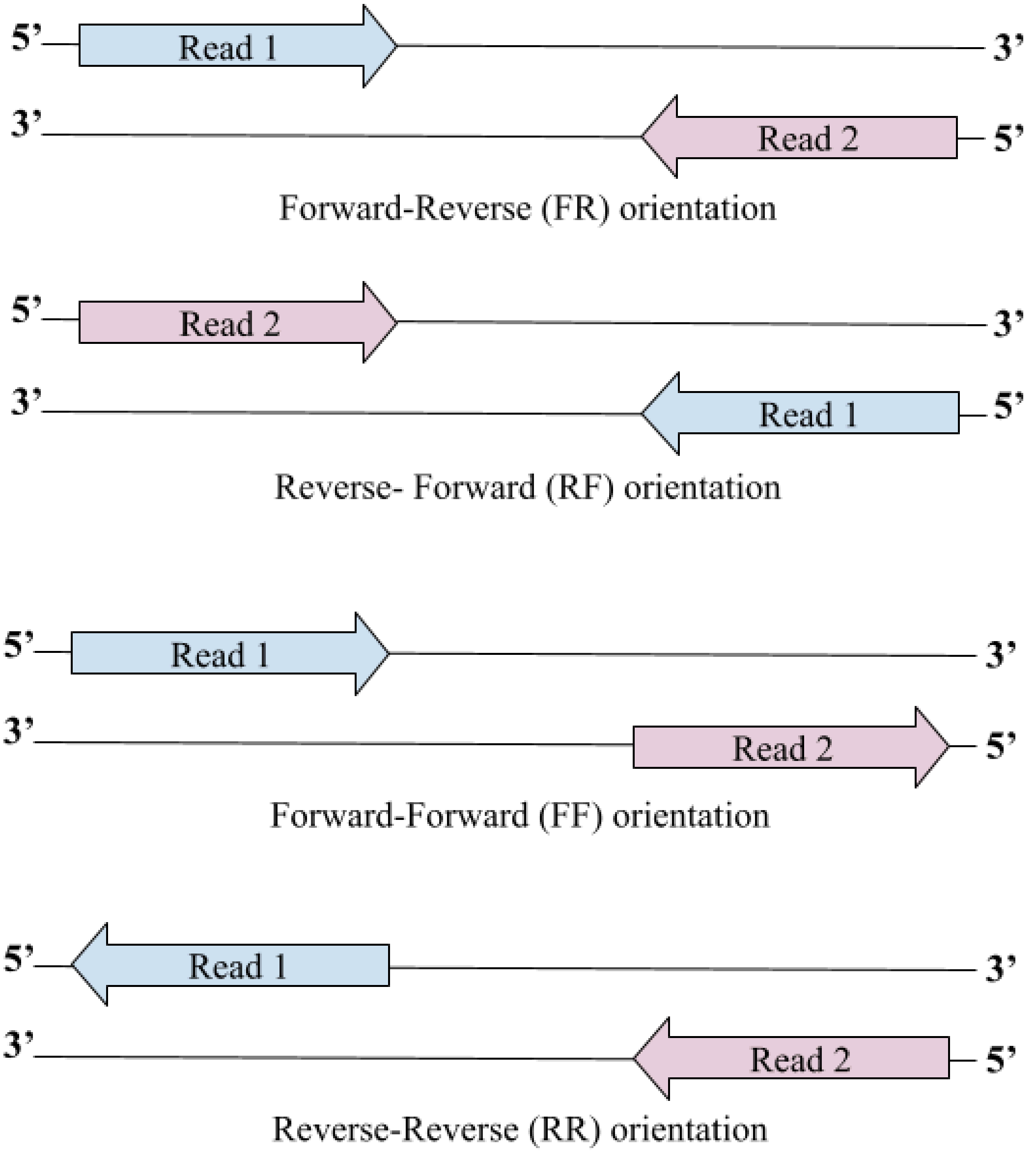

***Figure 3***: *Graphics showing different paired read orientations: the forward-reverse (FR) orientation, the reverse-forward (RF) orientation, the forward-forward (FF) orientation and the reverse-reverse (RR) orientation.*

### 5. Supplementary Results

#### 5.1. Impact of different model tuning

To evaluate the impact of tuning various models, we performed simulations of human genome sequencing in which specific model parameters were altered while all other user-defined parameters remained constant. The results of these simulations are presented below.

##### 5.1.1. MDA

Users can choose between a uniform amplification and MDA-like amplification with the '*coverage_distribution*' parameter. To check how these options change the coverage distribution of the sequencing simulation, we performed two simulations of the ScWGS approach of one cell each with a total read coverage of 0.03X. The simulated reads were then aligned to the reference genome using the Bowtie 2 (Langmead and Salzberg 2012) aligner. The aligned reads were sorted and the coverage for chromosome 1 was extracted from the data using Samtools (Li *et al.* 2009). The coverage distribution on chromosome 1 was plotted using a Python script as shown in Figure 4.

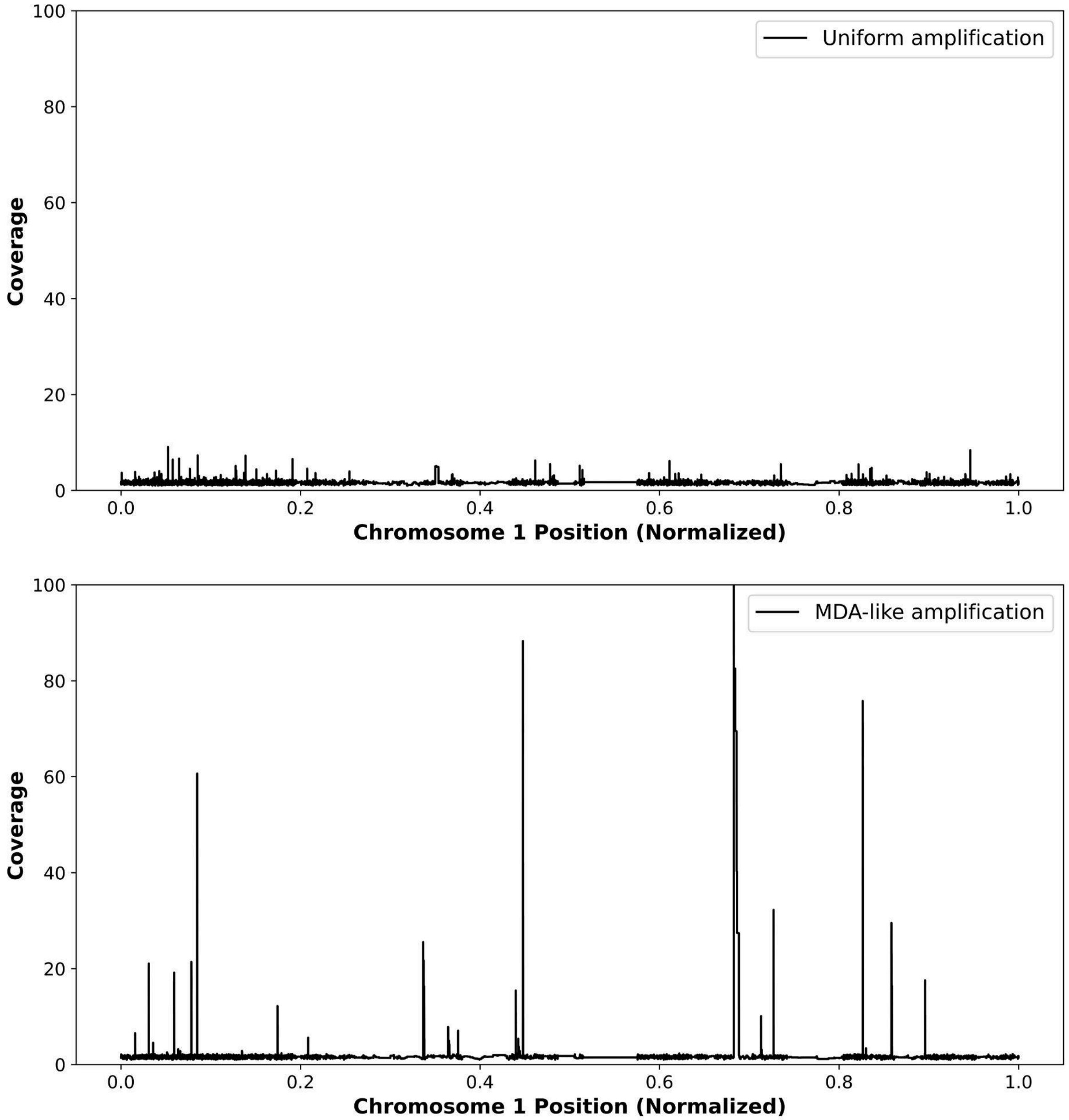

***Figure 4:*** *The read coverage distribution for chromosome 1 for the same coverage obtained from two simulations: (i) with a uniform amplification model (Top) and (ii) with a MDA-like amplification model (Bottom) are presented together for comparison.*

From the top plot in Figure 4, although we observe some regions are more amplified (small spikes) due to the GC bias across the genome, we see that the read coverage is mostly uniform at 0.03X across the chromosome when the amplification model is set to have uniform amplification. However, the bottom plot shows how this uniformity gets affected if the amplification model is changed to the classic MDA-like amplification approach that is expected to over-amplify certain genomic regions in a highly non-uniform manner (Gonzalez-Pena *et al.* 2021).

5.1.2. Beta Function

The Beta function describing the DNA fragmentation distribution is controlled by four user parameters: '*min_DNA_fragment_length*', '*max_DNA_fragment_length*', '*mode_DNA_fragment_length*', and '*beta_of_beta_distribution*'. These parameters respectively determine the minimum, maximum, mode, and beta values of the Beta distribution. The minimum and maximum values define the range of generated DNA fragment lengths, while the mode and beta values shape the fragment size distribution. Figure 5 illustrates the effect of different beta values on the distribution, given a minimum fragment length of 150 bp, a maximum fragment length of 1000 bp, and a mode of 350 bp.

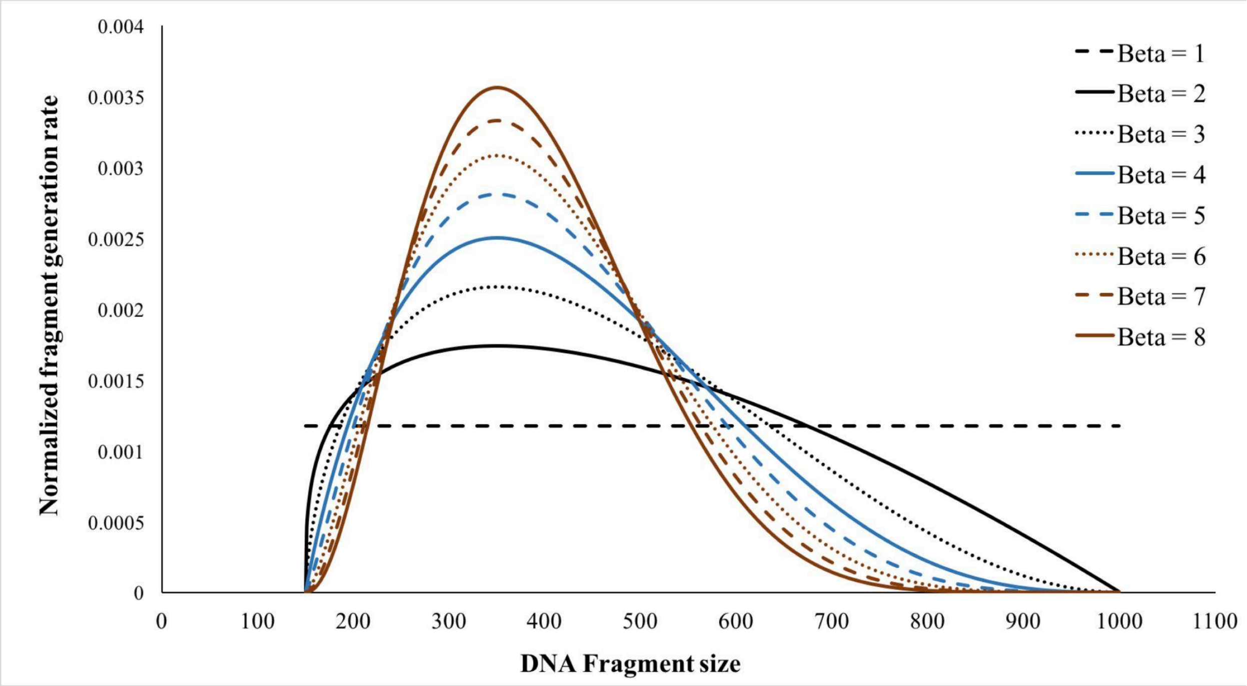

***Figure 5***: *The effect of varying the beta value in the Beta function describing the DNA fragment size distribution is shown for different beta values. The minimum DNA fragment length, maximum fragment length, and mode fragment length are set to 150 bp, 1000 bp, and 350 bp, respectively.*

5.1.3. GC Bias

In RadiSeq, GC bias is modeled using a triangle function. A GC bias value of 1.0 for all bins indicates that all genome regions have an equal likelihood of being sequenced, while a value of 0 means a region will not be sequenced. Users can control the tightness or looseness of this bias by setting the '*degree_of_GC_bias*' parameter, which determines the slope of the triangle function. The GC fraction (the ratio of guanine and cytosine bases to the total bases) will then determine the GC bias for each region based on the selected degree of bias. Table 2 shows the GC fraction ranges for which the GC bias is non-zero, across different degrees of GC bias.

***Table 2****: The range of GC fraction for which the GC bias is non-zero is shown for different degrees of GC bias.*

| Degree of GC bias | GC fraction range with non-zero bias |
|---|---|
| 0 | 0.0 - 1.0 |
| 0.05 | 0.3 - 0.7 |
| 0.1 | 0.4 - 0.6 |
| 0.2 | 0.45 - 0.55 |
| 0.4 | 0.475 - 0.525 |
| 1.0 | 0.49 - 0.51 |

To characterize the effect of this bias on the read coverage, two BcWGS simulations were performed with uniform amplification and a total coverage of 0.03X for a single cell. One of the simulations used a degree of GC bias value of zero (unbiased – without GC bias) and the other used a value of 0.1 (biased – with GC bias). The generated reads were aligned using Bowtie 2 aligner and the read coverage for chromosome 1 was extracted using the Samtools.

Figure 6 compares the read coverage distribution on chromosome 1 from the unbiased and biased simulations. When the GC bias is set to zero, the coverage is very uniform as expected and it is equally likely that the reads can get generated anywhere from the chromosome as shown in the top plot in Figure 6. With the GC bias on the other hand, we observed that the uniformity in coverage is modified according to the GC content of each genomic region. In particular, we can see that the region in the middle of chromosome 1, which corresponds to the centromere, gets a very low coverage. This is expected since the centromere is observed to be a GC-poor trough (Lynch *et al.* 2010).

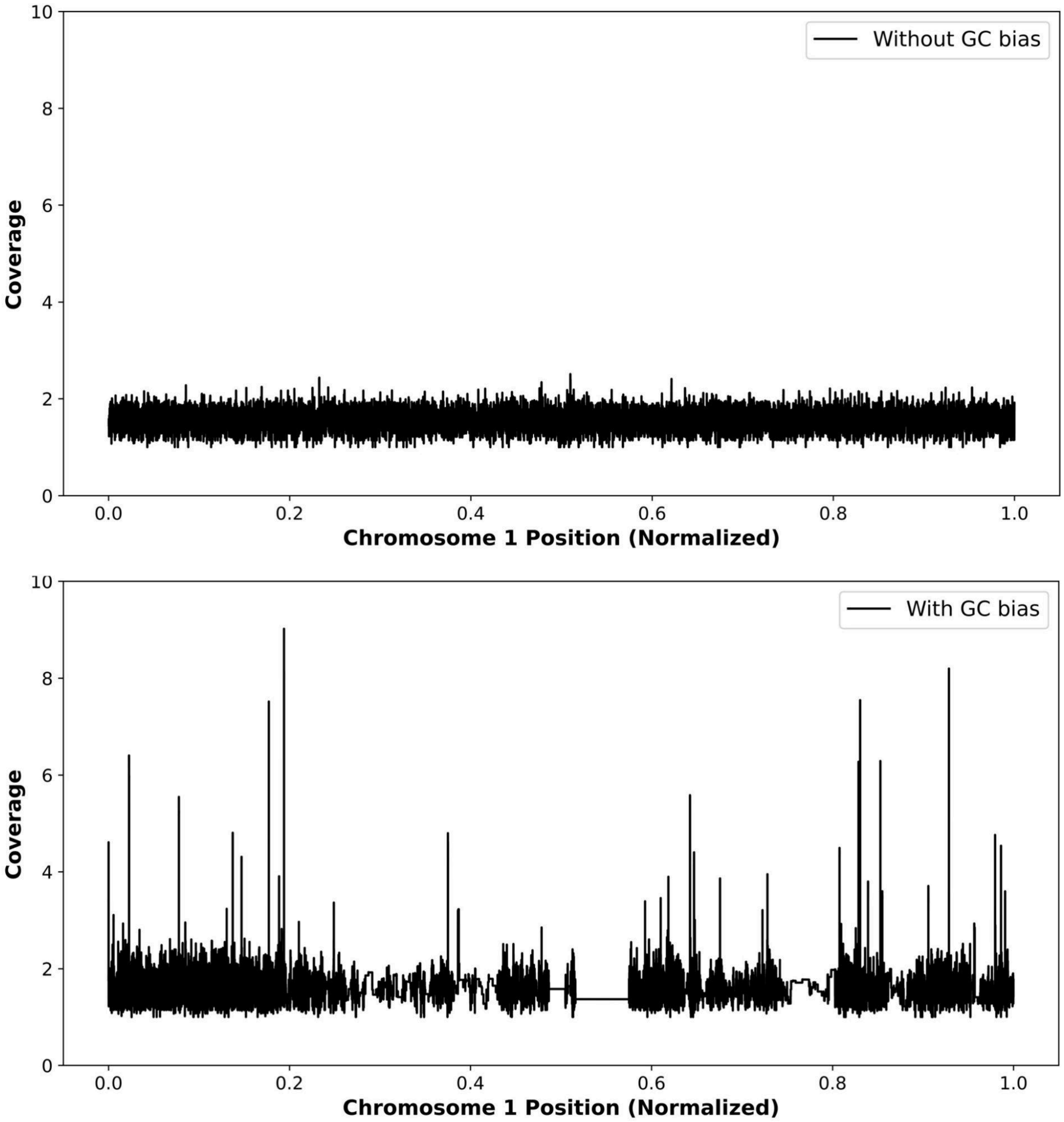

***Figure 6****: The read coverage distribution for chromosome 1 obtained from two simulations: (i) with zero GC bias (Top) and (ii) with a degree of GC bias value of 0.1 (Bottom) are presented together for comparison.*

5.1.4. Chimeric Read

In RadiSeq, users can control the rate of chimera formation with the '*read_artifacts_rate*' parameter. To characterize the effect of this parameter on the generated read data, BcWGS simulations were repeated with uniform amplification and a total coverage of 0.03X but with varying read artifact rates. The simulated reads were aligned to the reference genome using the Bowtie 2 aligner and the aligned data was analyzed using Samtools. In the Samtools summary statistics, we observed that the read artifacts rate was affecting the number of reads mapped and the reads paired significantly. Figure 7 shows the impact of varying read artifacts rates on the number of reads mapped and the number of reads properly paired out of all the generated reads.

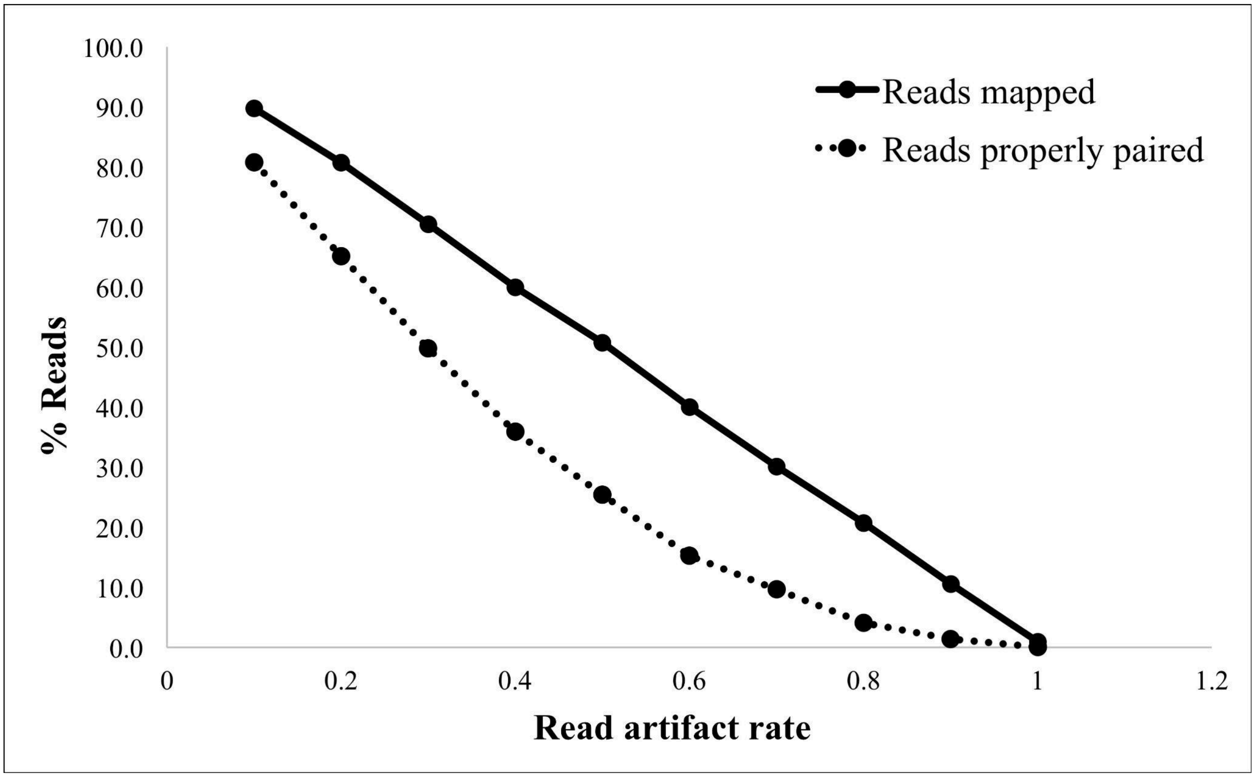

***Figure** 7: Graph showing the change in the number of reads mapped and the reads properly paired with the change in read artifact rate in RadiSeq simulation.*

5.1.5. Read Orientation

RadiSeq includes the option to simulate different read orientations in paired-end read data. Repeated BcWGS simulations were conducted with varying '*fraction_of_other_oriented_read_pairs*' at a coverage of 0.03X while keeping all other parameters constant. The reads were aligned with Bowtie 2 and analyzed using Samtools. The summary statistics were then compared across simulations to characterize the impact of the parameter for the fraction of other-oriented read pairs in RadiSeq.

Figure 8 shows the change in the number of read pairs in forward-reverse (FR orientation), reverse-forward (RF orientation) and other read pair orientations with the change in the fraction of other-oriented read pairs parameter in RadiSeq. As expected, with the increase in the fraction of other-oriented read pairs, the percentage of FR read pairs (inward oriented) read pairs decreased linearly, while the percentages of both the RF read pairs (outward oriented) and read pairs in other orientations increased.

Interestingly, the change in read pair orientation also impacted the read mapping quality as well as the read pairing efficiency. As shown in Figure 9, the number of reads mapped increased as the fraction of other-oriented read pairs increased. However, the increase in the fraction of other-oriented read pairs inversely affected the number of reads that are properly paired. Additionally, we have also observed that the change in the number of reads mapped and reads properly paired is also correlated with the number of read pairs on different chromosomes and the mean insert size in the sample respectively.

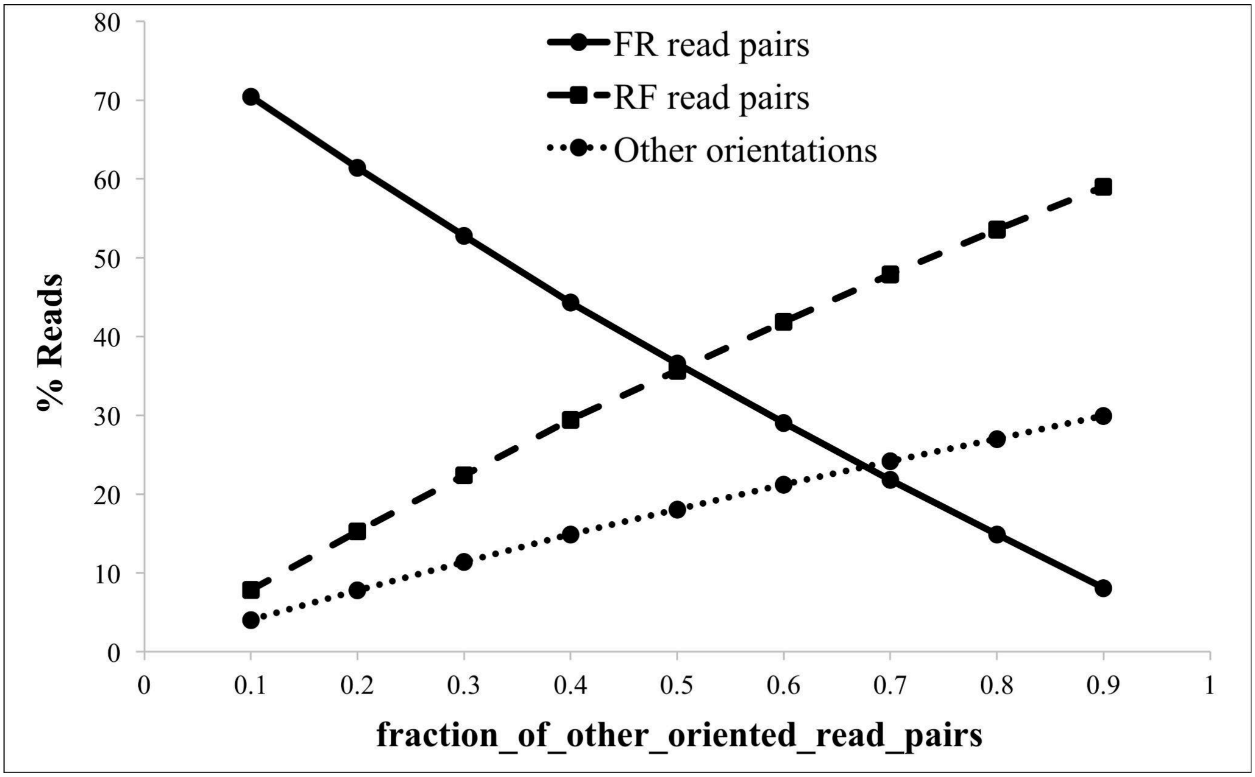

***Figure 8***: *Graph showing the change in the number of read pairs in different read orientations with the change in the 'fraction_of_other_oriented_read_pairs' parameter in RadiSeq.*

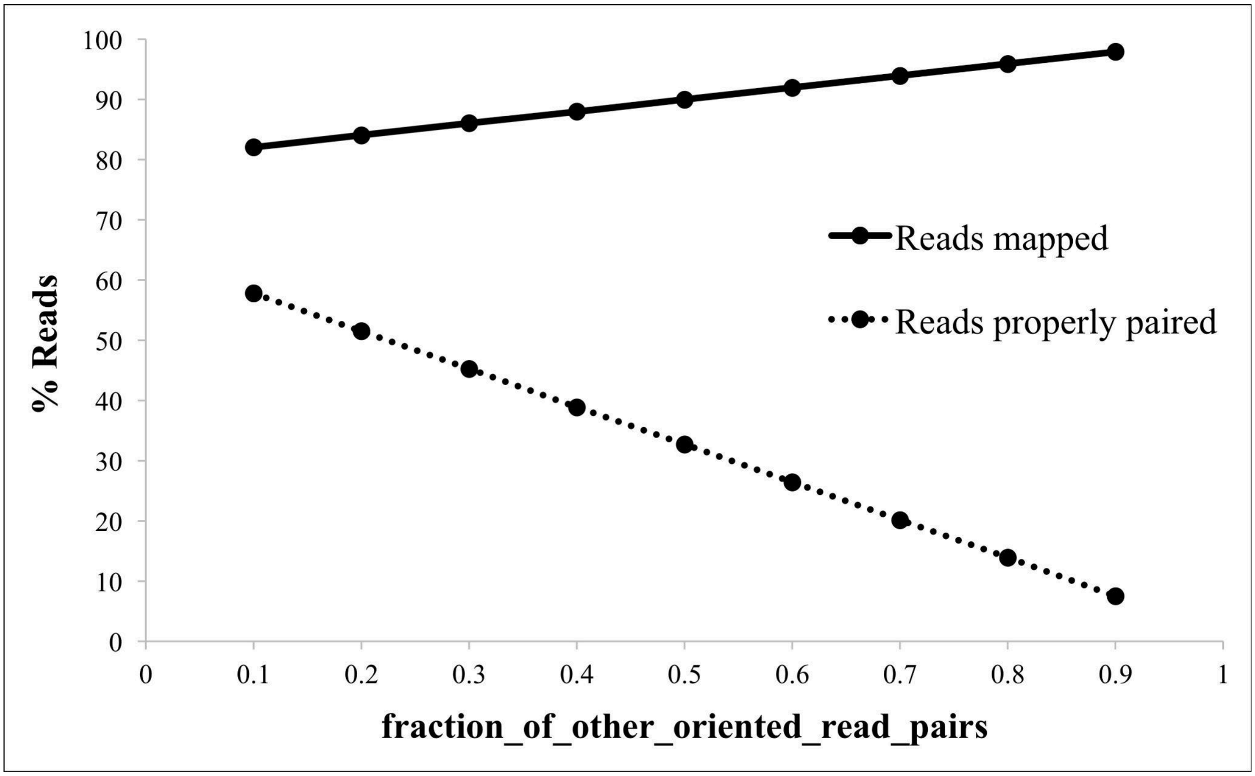

***Figure 9***: *Graph showing the change in the number of reads mapped and the reads properly paired with the change in the 'fraction_of_other_oriented_read_pairs' parameter in RadiSeq.*

### 5.2. FastQC results

The FastQC tool (http://www.bioinformatics.babraham.ac.uk/projects/fastqc/) was used to assess the RadiSeq-generated reads. The FastQC results for the RadiSeq data were compared to those obtained from the ScWGS experimental data that we had. Overall, a reasonable agreement was observed between the RadiSeq simulated data and the experimental data. Figures 10, 11, and 12 show the FastQC-generated plots of 'Per Base Sequence Quality', 'Sequence Length Distribution', and 'Sequence Duplication Levels', respectively, for one cell sequenced using the ScWGS technique both computationally and experimentally.

5.2.1. <u>Per Base Sequence Quality Plot</u>

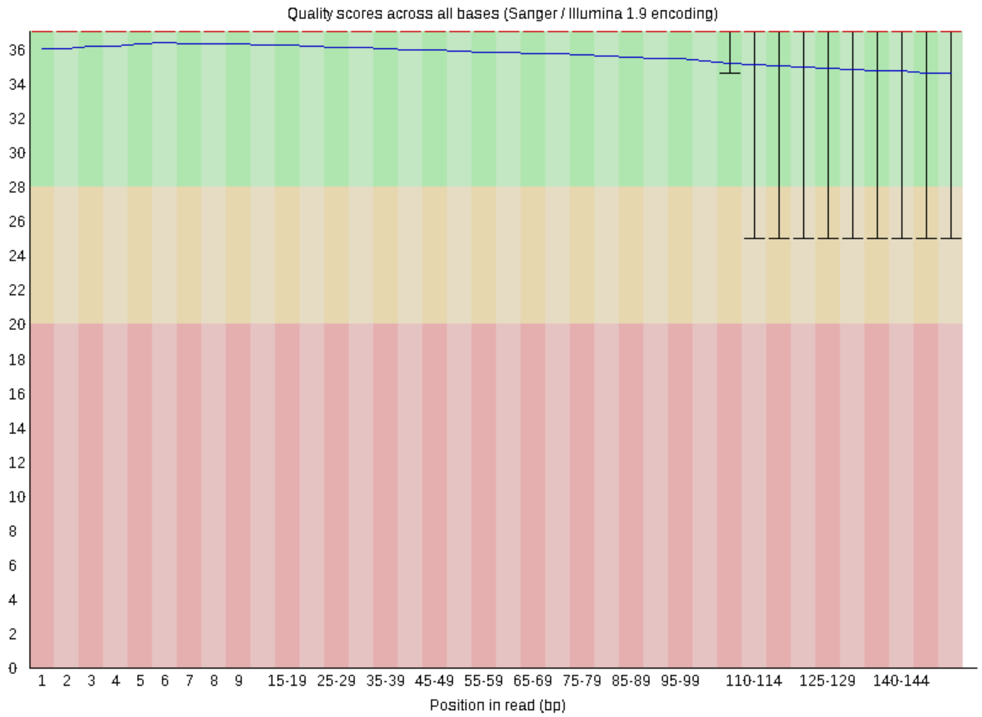

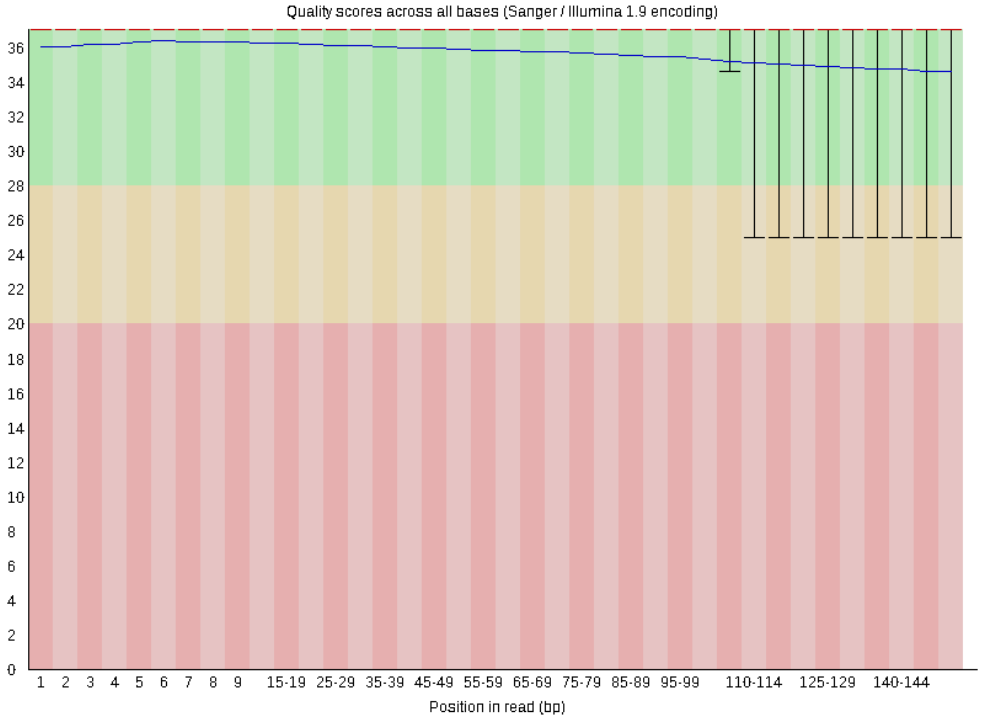

***Figure 10***: *The Phred quality score distribution across the different read positions is shown in the FastQC generated 'per base sequence quality' plots. The experimentally obtained read data (Top) and RadiSeq-generated data (Bottom) are compared in this figure. The Y-axis is showing the Phred score for each location in the read on the X-axis.*

5.2.2. <u>Sequence Length Distribution Plot</u>

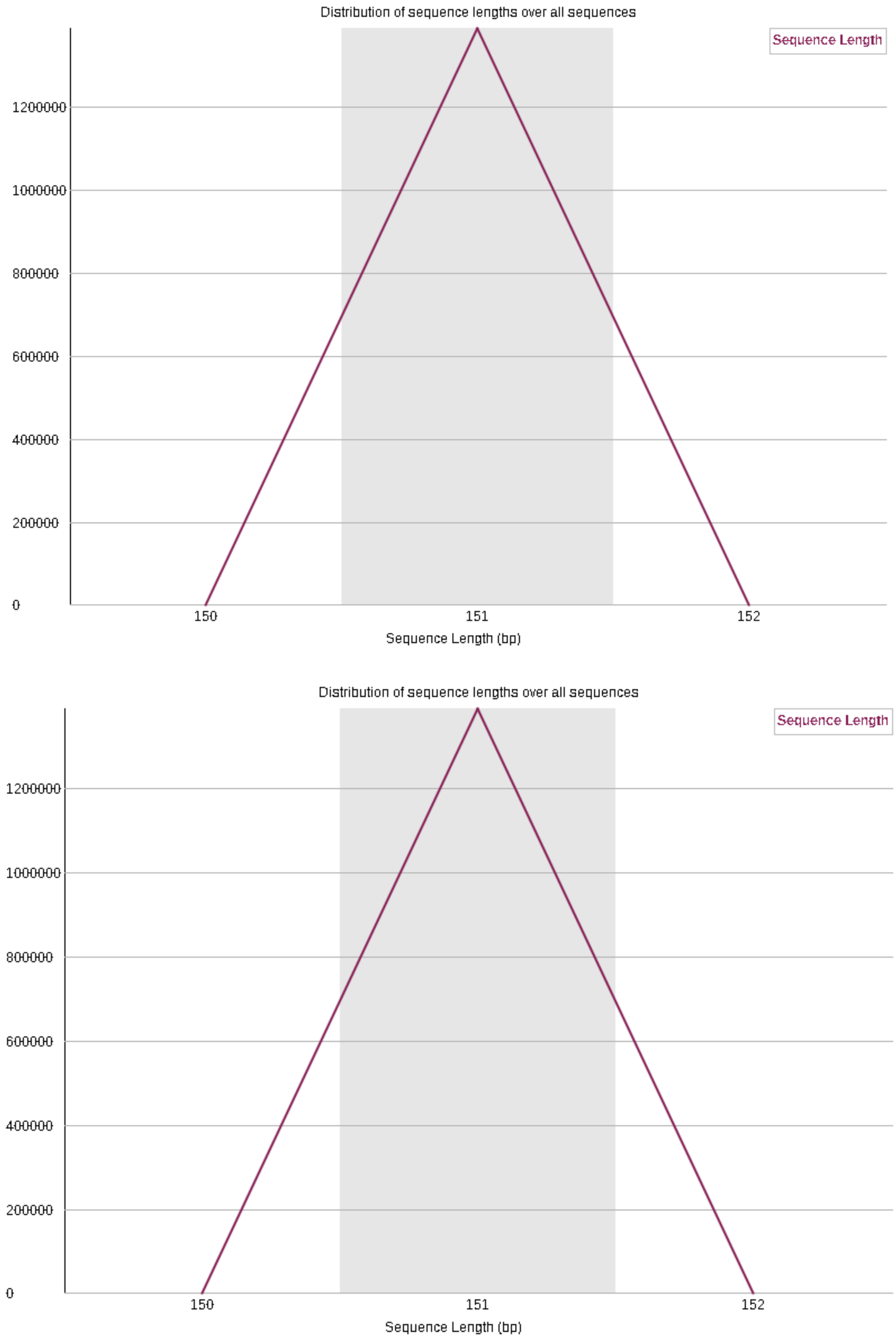

***Figure 11****: The distribution of the read sequence lengths for the sample is shown in the FastQC generated 'sequence length distribution' plots. The experimentally obtained read data (Top) and RadiSeq-generated data (Bottom) are compared in this figure. The Y-axis is showing the number of read sequences with a specific length given on the X-axis.*

5.2.3. <u>Sequence Duplication Levels Plot</u>

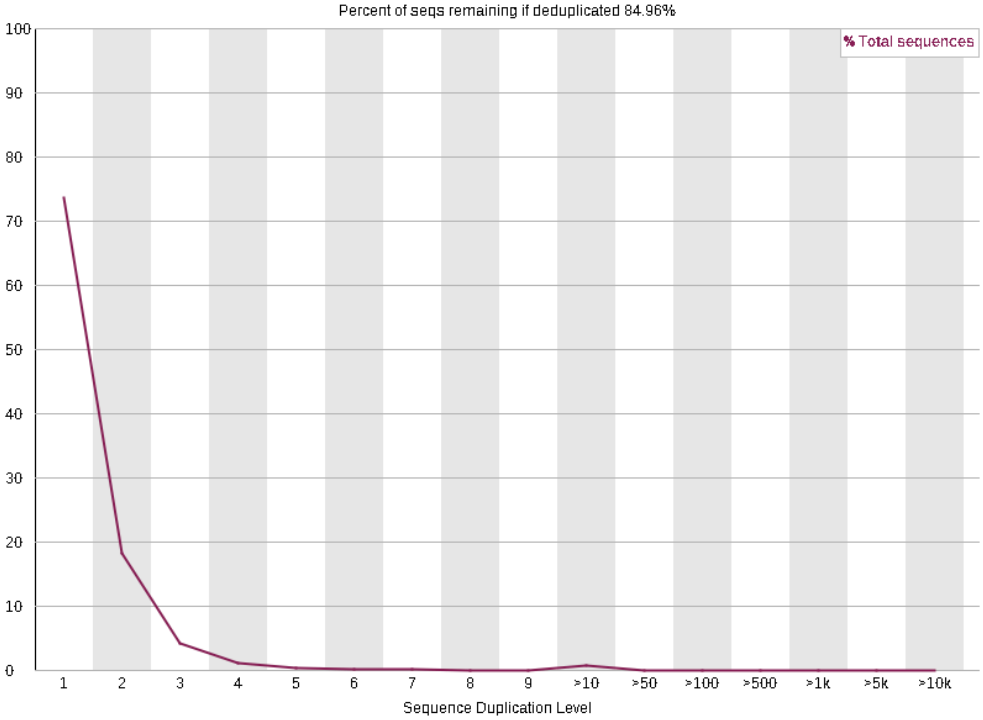

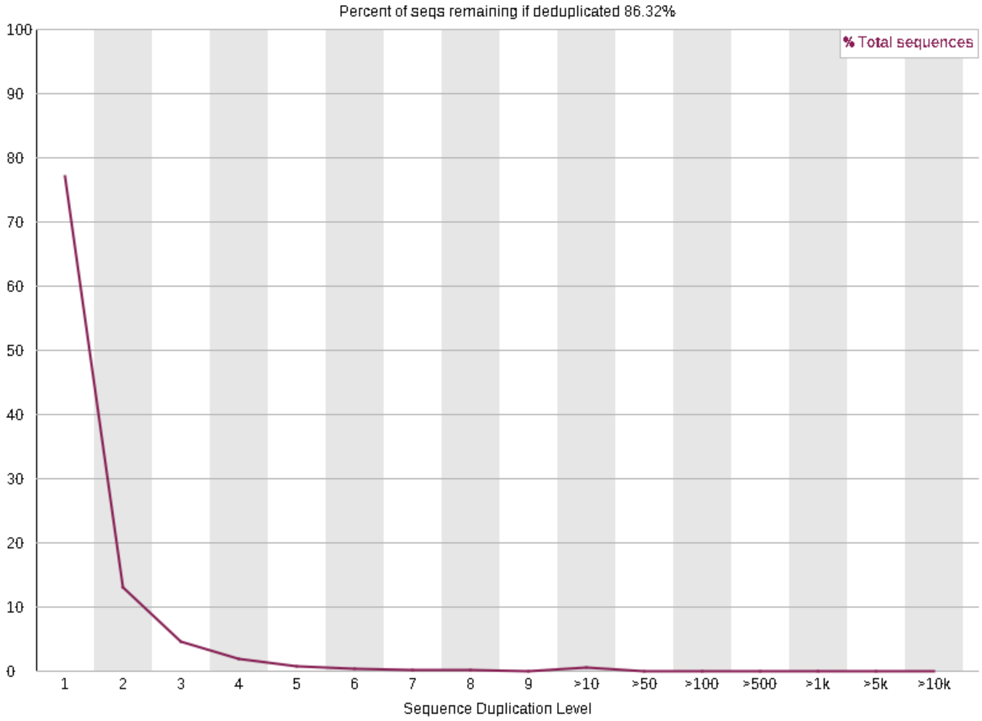

***Figure 12****: The percentage of sequence duplication is shown in the FastQC generated 'sequence duplication levels' plots. The experimentally obtained read data (Top) and RadiSeq-generated data (Bottom) are compared in this figure. The Y-axis is showing the percentage of sequences that are duplicated for a specific duplication level given on the X-axis.*

### 5.3. Simulation accuracy

To validate the simulation accuracy of RadiSeq, simulated data were compared against the experimental data. Experiments were performed on a human B-lymphoblastoid cell line that we cultured in our lab. These cells belonged to a well-characterized genome of an Ashkenazi individual (Shumate *et al.* 2020). Radiation-exposed and sham-irradiated cells were sequenced both using ScWGS and BcWGS techniques using Illumina NovaSeq 6000 sequencer. We used the sham-irradiated control data to compare against the RadiSeq-generated sequence data.

For RadiSeq simulations, the input parameters were made to match the experimental conditions and the data were aligned and processed identically as with the experimental data. Using Samtools, the aligned data were analyzed and the key statistics were compared between experiment and simulation to check for discrepancies and similarities. A good agreement of the RadiSeq-generated data with the experiment confirmed the accuracy of our simulations.

Figure 13 compares the read mapping quality score distributions obtained after the BcWGS experiment and simulations. Figure 14 compares the same distributions but for ScWGS.

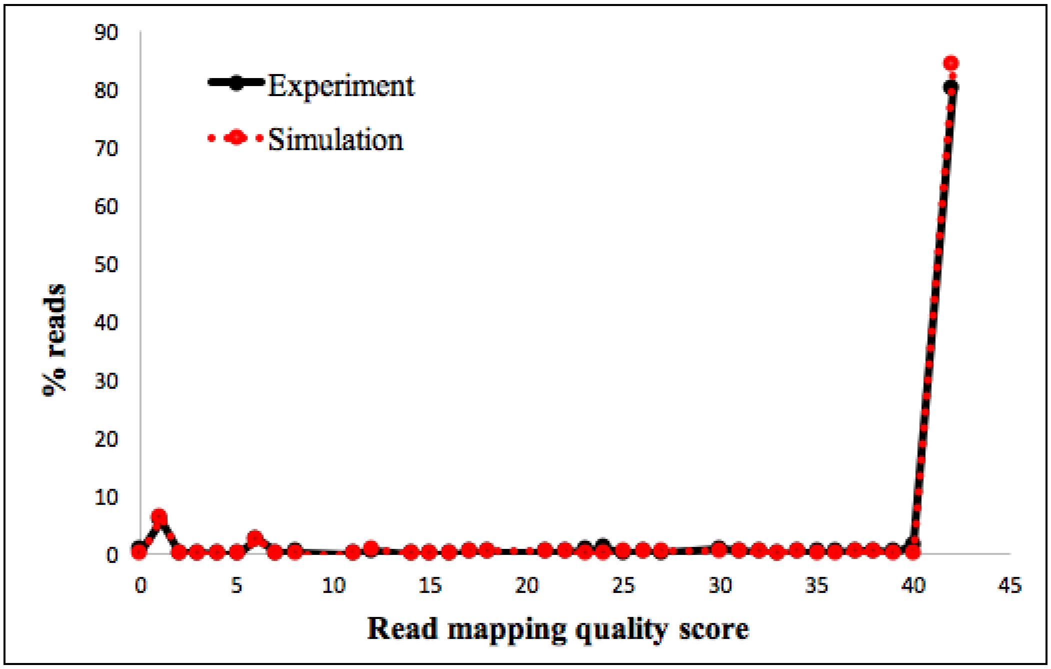

***Figure 13**: The graph comparing the read mapping quality score distribution of the experimentally obtained (black) and RadiSeq simulated (red) reads data from bulk-cell whole genome sequencing.*

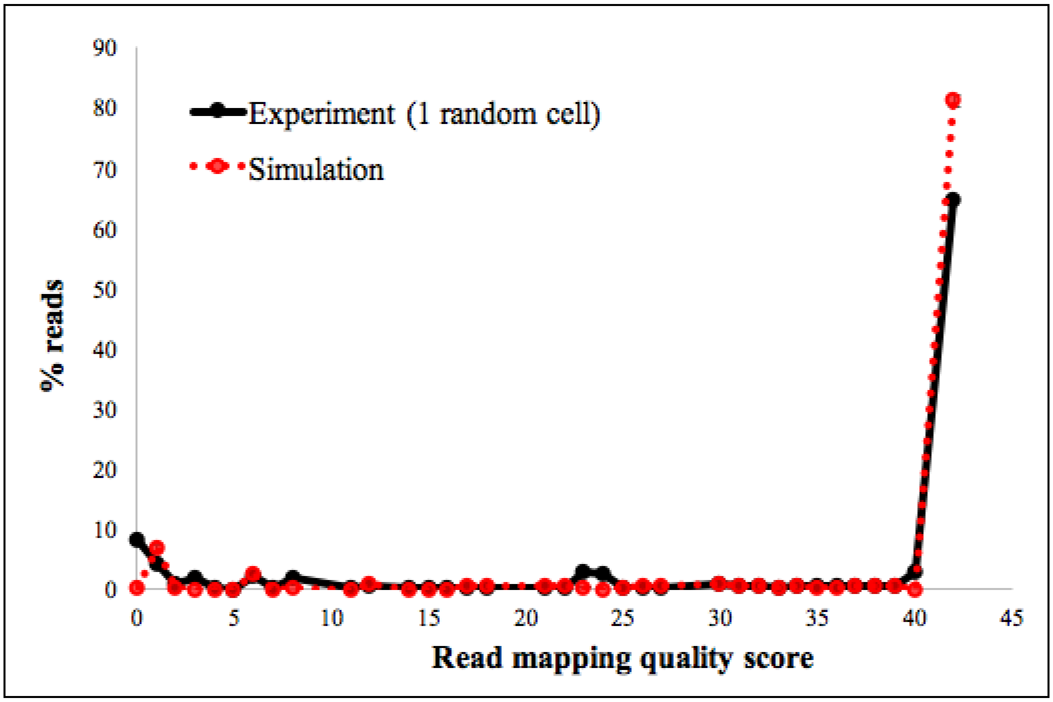

***Figure 14**: The graph comparing the read mapping quality score distribution of the experimentally obtained and RadiSeq simulated reads data from single-cell whole genome sequencing. Two single cells are randomly picked from the experiment to compare against the RadiSeq simulated data.*

### 6. References

Benjamini Y, Speed TP. Summarizing and correcting the GC content bias in high-throughput sequencing. *Nucleic Acids Res* 2012;**40**:e72.

Chen Y-C, Liu T, Yu C-H *et al.* Effects of GC bias in next-generation-sequencing data on de novo genome assembly. *PLoS One* 2013;**8**:e62856.

Diossy M, Sztupinszki Z, Krzystanek M *et al.* Strand Orientation Bias Detector to determine the probability of FFPE sequencing artifacts. *Brief Bioinform* 2021;**22**, DOI: 10.1093/bib/bbab186.

Erlich HA, Gibbs RA, Gibbs R *et al. Polymerase Chain Reaction.*, 1989.

Gonzalez-Pena V, Natarajan S, Xia Y *et al.* Accurate genomic variant detection in single cells with primary template-directed amplification. *Proc Natl Acad Sci U S A* 2021;**118**, DOI: 10.1073/pnas.2024176118.

Gupta AK, Nadarajah S. *Handbook of Beta Distribution and Its Applications*. CRC Press, 2004.

Huang W, Li L, Myers JR *et al.* ART: a next-generation sequencing read simulator. *Bioinformatics* 2012;**28**:593–4.

Langmead B, Salzberg SL. Fast gapped-read alignment with Bowtie 2. *Nat Methods* 2012;**9**:357–9.

Li H, Handsaker B, Wysoker A *et al.* The Sequence Alignment/Map format and SAMtools. *Bioinformatics* 2009;**25**:2078–9.

Lu N, Qiao Y, Lu Z *et al.* Chimera: The spoiler in multiple displacement amplification. *Comput Struct Biotechnol J* 2023;**21**:1688–96.

Lynch DB, Logue ME, Butler G *et al.* Chromosomal G + C content evolution in yeasts: systematic interspecies differences, and GC-poor troughs at centromeres. *Genome Biol Evol* 2010;**2**:572–83.

Mathew F, Kildea J. *kildealab/RadiSeq: RadiSeq_v2.0*. Zenodo, 2024.

Matsumoto M, Nishimura T. Mersenne twister. *ACM Trans Model Comput Simul* 1998;**8**:3–30.

Montgomery L, Lund CM, Landry A *et al.* Towards the characterization of neutron carcinogenesis through direct action simulations of clustered DNA damage. *Phys Med Biol* 2021;**66**, DOI: 10.1088/1361-6560/ac2998.

Schneider VA, Graves-Lindsay T, Howe K *et al.* Evaluation of GRCh38 and de novo haploid genome assemblies demonstrates the enduring quality of the reference assembly. *Genome Res* 2017;**27**:849–64.

Shumate A, Zimin AV, Sherman RM *et al.* Assembly and annotation of an Ashkenazi human reference genome. *Genome Biol* 2020;**21**:129.

Spits C, Le Caignec C, De Rycke M *et al.* Whole-genome multiple displacement amplification from single cells. *Nat Protoc* 2006;**1**:1965–70.

Thakur J, Packiaraj J, Henikoff S. Sequence, Chromatin and Evolution of Satellite DNA. *Int J Mol Sci* 2021;**22**, DOI: 10.3390/ijms22094309.
